# Spatial architecture of complex antibody formats dictates pharmacokinetics via FcRn-recycling

**DOI:** 10.64898/2026.11.02.763654

**Authors:** Stefan C. Weise, Cordula Jany, Sabine Imhof-Jung, Claire Simonneau, Johannes Fraidling, Anna-Lena Bolender, Aman Padamsey, Elisa Di Lenarda, Anne Christine Cascais, Thomas Emrich, Tilman Schlothauer, Annette Indlekofer, Hubert Kettenberger, Volker Haucke, Wolfgang F. Richter, Thomas E. Kraft

## Abstract

Complex antibody formats are advancing in the biotherapeutic landscape to address the need for multi-specificity, tailored valency, and novel mechanisms of action. However, understanding how non-canonical architectures impact fundamental pharmacokinetic pathways remains a critical challenge. While canonical IgGs experience steric hindrance at the endosomal membrane during FcRn engagement, whether – and how strongly – this constraint impacts FcRn-mediated recycling across the expanding diversity of non-canonical antibody architectures has remained unknown. By systematically evaluating a diverse panel of rationally designed antibody geometries utilizing orthogonal in vitro assays we demonstrate that spatial layout profoundly influences FcRn-recycling efficiency; inverting the antibody structure by positioning the Fab-domains to the C-terminus (‘invertbodies’) maximizes recycling, whereas crowding the N-terminus severely impairs it. In contrast, non-specific endocytosis was only modestly and inconsistently affected by Fab number, dependent on the Fv-sequence, but not by the spatial architecture itself. Notably, antibody architecture and affinity-enhancing Fc-FcRn mutations act independently yet cooperatively, ultimately revealing a physiological ‘recycling ceiling’. These FcRn-recycling dynamics translate with high fidelity to in vivo clearance (CL) in a humanized FcRn mouse model. Across the antibody panel, architecture drove a near 15-fold variance in CL, an effect that collapsed in FcRn-knockout mice. As proof of concept, we applied the invertbody design to a variant of the marketed antibody golimumab, achieving a three-fold reduction in clearance without compromising potency. These findings reveal the magnitude and predictability with which the antibody architecture governs FcRn-recycling and non-specific CL, establishing a rational, generalizable framework to optimize the pharmacokinetics of biotherapeutics.

## INTRODUCTION

In the past, canonical, bivalent IgGs constituted the majority of approved therapeutic antibodies. However, advancements in antibody engineering technologies have paved the way for the generation of complex antibody formats (1–3). Currently, 112 such formats are in clinical development (4). These novel architectures, which include an array of multivalent and multispecific antibodies, offer the potential to engage multiple targets simultaneously, enhance therapeutic efficacy, and address complex, unmet medical needs with superior flexibility (1–3). Yet, despite the clinical advantage, there is currently no comprehensive understanding of how these complex formats impact fundamental pharmacokinetic (PK) properties. This knowledge gap poses the risk of highly unpredictable clearance rates, resulting in lengthy and costly optimization cycles during preclinical development or undesirable PK properties of the clinical molecule.

Favorable PK, particularly low systemic clearance or high bioavailability after subcutaneous administration, is critical for managing chronic diseases. Optimizing these parameters allows for less frequent dosing, which directly improves patient convenience, enhances clinical adherence and represents an essential factor of therapeutic success (5–8). Consequently, the early elimination of candidates with unfavorable PK profiles and the rational optimization of molecules with superior exposure kinetics are paramount during the discovery and preclinical phases (9, 10).

Systemic clearance (CL) is the primary PK parameter quantifying the rate of antibody elimination from the body, mediated by a combination of target-specific and target independent biological processes. While target-mediated CL pathways are inherently saturable and restricted to specific drug-target pairs, non-specific CL represents the continuous, baseline elimination pathways that universally impact all immunoglobulin G (IgG) antibodies (11).

This non-specific CL is driven by non-specific endocytosis across various tissues, followed by intracellular sorting for either lysosomal degradation or FcRn-mediated salvage (12–15). Constitutive fluid phase pinocytosis, which captures bulk extracellular fluid non-selectively (16, 17), and adsorptive endocytosis (18) mediate the cellular internalization. In the latter, electrostatic interactions between positively charged antibody surfaces, particularly within the variable domains, and the anionic components of the glycocalyx enrich local concentration on the cell surface, resulting in increased cellular uptake (19, 20). This charge-mediated uptake mechanism represents a significant liability, as it markedly increases cellular internalization, leading to accelerated antibody clearance and consequently, reduced in *vi*v*o* exposure and efficacy (18, 19, 21, 22).

Following internalization, the interaction with the neonatal Fc receptor (FcRn) within acidified endosomes determines the intracellular fate of an IgG. FcRn is a membrane-bound, heterodimeric receptor primarily located within endosomes, composed of a ꞵ2-microglobulin subunit and a FcRn-alpha chain. At the lowered endosomal pH, two FcRn molecules can form a complex with one IgG by binding the Fc region, which initiates sorting it into recycling endosomes for exocytosis back to the pH neutral extracellular environment. Unbound antibodies or proteins are directed to late endosomes for lysosomal elimination (23–25). Notably, FcRn employs an analogous pH-dependent mechanism to salvage endocytosed serum albumin via a distinct, non-overlapping FcRn binding site, extending the half-life of both major long-lived serum proteins (25).

In clinical settings, therapeutic antibodies (typically with C_max_ in the range between 10 - 400 ug/mL (9)) must compete for a finite amount of endosomal FcRn against a 50 - 1000 fold excess of endogenous IgGs (approximately 10 - 16 mg/mL (26)). To overcome this competition and improve antibody salvage, half-life extension strategies rely on affinity engineering of the antibody’s Fc domain. These include for example the YTE (27), LS (8) and ACT5 (28) mutations, which enhance pH-dependent FcRn binding and markedly extend serum half-life in approved biotherapeutics, such as Nirsevimab and Sotrovimab (29).

Structural models of canonical IgGs bound to FcRn indicate that the Fab domains in these Y-shaped IgGs affect FcRn engagement (30, 31). Because the Fc-domain binds to FcRn in an ‘upside-down’ orientation, the bulky Fab-domains project directly towards the lipid-bilayer of the endosomal vesicle. This physical alignment introduces severe spatial restrictions, forcing the Fab domains of the bound IgG to adopt a more compact, T-shaped or reclined conformation to minimize physical crowding against the membrane boundary (25, 30, 31). Because this membrane-dependent steric hindrance relies strictly on the proximity to a physical bilayer, its effect remains hidden in cell-free assays, such as surface plasmon resonance, while manifesting clearly in cell-based assays, as demonstrated for whole IgG compared to Fc-only molecules (31). While these structural models describe steric constraints for canonical IgGs, whether and with what consequences, this constraint extends to the spatial architecture of non-canonical, complex antibody formats remains unknown.

Recapitulating the dual-step process (endocytosis and recycling) within a matching molecular environment to predict non-specific CL properties requires a relevant cellular model. Because liver sinusoidal endothelial cells (LSECs) act as key players in non-specific CL due to their high endocytic activity, abundant glycocalyx coverage, elevated exposure based on their anatomical position and abundant FcRn-expression (12–15), they represent an ideal physiological cell type for developing membrane-competent in vitro cellular assays.

While in vitro assays are valuable tools to isolate endocytosis and FcRn-recycling mechanisms, quantitative assessment of systemic non-specific CL still requires predictive in vivo models. Because wild-type mouse FcRn has an increased affinity for human IgG compared with human FcRn, transgenic human FcRn Tg32 mice (mFcRn-/-, hFcRn Tg32) are preferred as a model to estimate human antibody clearance. In these mice, the endogenous murine FcRn is knocked out and replaced by the human FcRn alpha chain, which results in a more physiological binding of human IgG (32–34). However, since human FcRn binds endogenous murine IgG poorly, these mice exhibit abnormally low baseline levels of endogenous murine IgG due to accelerated CL (35). To account for the (50-1000 fold) excess of circulating, endogenous antibodies present in patients, high-dose intravenous human immunoglobulins (IVIG) can be co-administered with the therapeutic antibody to simulate a competitive endosomal binding environment for FcRn. These pooled human IgG antibodies provide a physiological model of FcRn saturation and recycling dynamics as seen in human biology (6, 36–38).

Here, we investigated the impact of three antibody format design principles on FcRn-recycling and non-specific clearance, using a diverse panel of rationally designed antibody architectures. By combining membrane-competent cellular LSEC assays with translational FcRn mouse models, we systematically decoupled binder-driven endocytosis from architecture-driven FcRn-recycling. Our results demonstrate that spatial antibody architecture predictably determines FcRn-recycling efficiency, while non-specific endocytosis remains comparatively architecture-independent. This architecture-dependent FcRn-recycling efficiency, in turn, is a highly predictable driver of non-specific CL, establishing architectural engineering as a powerful framework for the rational design of next-generation biotherapeutics.

## RESULTS

### 1. The spatial antibody architecture determines the efficiency of FcRn-recycling

Structural models of the membrane-anchored IgG:FcRn complex indicate that the antibody assumes a specific orientation in which the Fab arms are directed toward the endosomal membrane (30, 31) (Figure 1A). This spatial arrangement suggests physical constraints at the membrane interface that destabilize the complex (25, 30, 31), yet how these constraints impact FcRn-recycling efficiency and whether this scales with the spatial complexity of non-canonical antibody architectures has not been tested. To systematically investigate this, we designed a panel of 14 complex antibody formats (Figure S1A) based on three rational design rules: (I) increasing steric hindrance via additional N-terminal Fabs (Figure 1B and E, format-keys: N-N, N-NN, N-NNN, NN-NN), (II) avoiding hindrance by relocating Fabs to the C-terminus (Figure 1C and F, format-keys: C-C, C-CC, CC-CC), and (III) reducing steric constraints using one-Fab-armed configurations (Figure 1D and G, format-keys: 0-N, 0-NN, 0-NNN). An additional fourth group (IV) included formats with more unconventional (contorsbody (39)) or simultaneous N-& C-terminal arrangements, which did not directly match the design rules (Figure S1B, format-keys: cont-cont, 0-NC, N-NC, NC-NC). To isolate the effect of the Fab arrangement, we utilized consistent building blocks across the panel, including identical non-binding Fv domains (Binder#1), the same, effector function-less Fc-domain (PG-LALA), and consistent GS-linkers.

**Figure 1.**
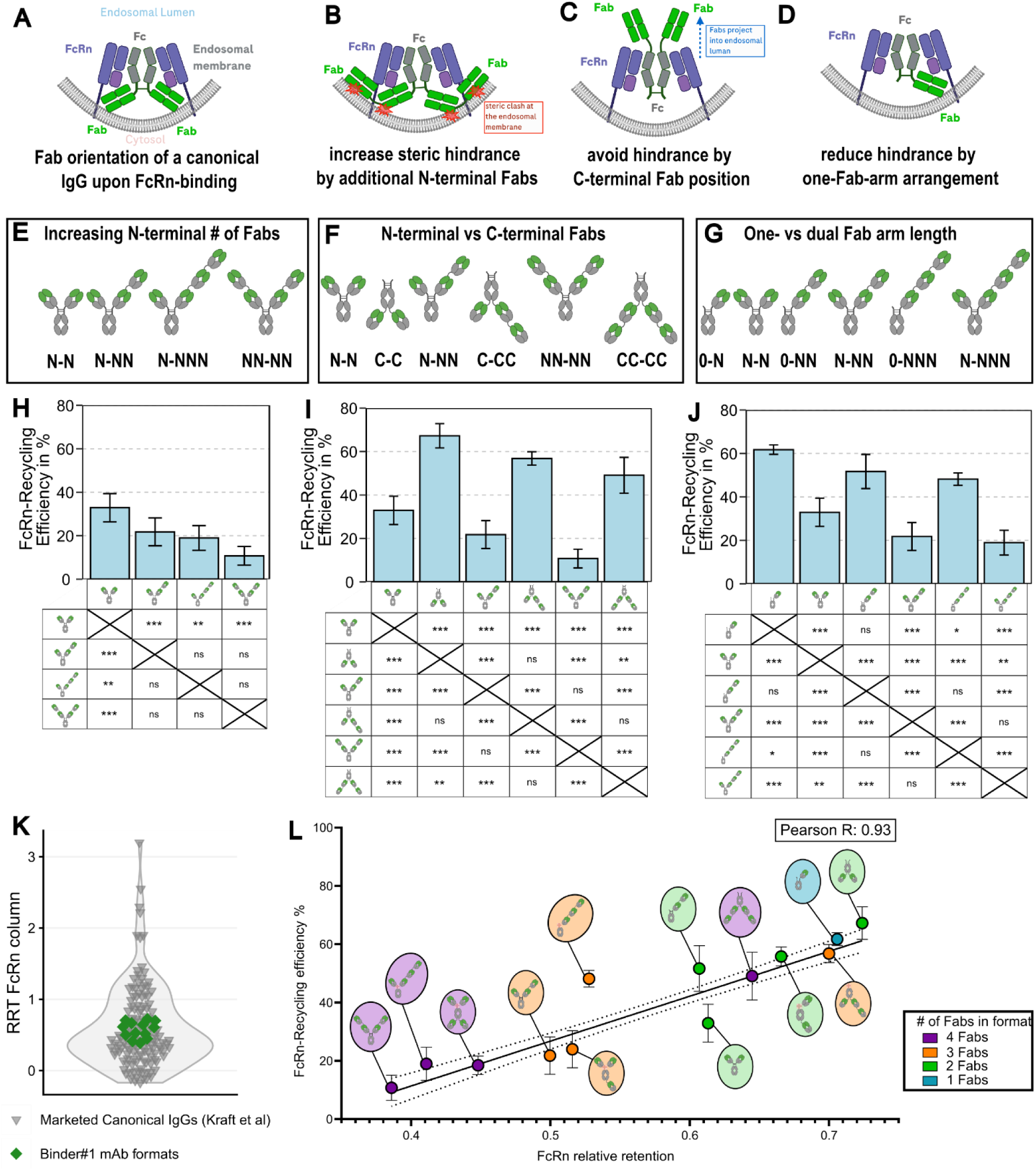
The complex antibody format impacts FcRn-recycling by architectural rules. (**A**) Schematic model of the interaction between a canonical IgG and the membrane bound FcRn demonstrating steric constraints at the endosomal membrane. (**B** – **D**) Schematic models illustrating the three hypothesized design principles for representative complex antibody formats. (**B**) Increasing the N-terminal number of Fabs increases clash at the endosomal membrane. (**C**) C-terminal Fabs reach into the endosomal lumen and avoid membrane interaction. (**D**) The one-armed antibody design reduces steric hindrance, facilitating Fab arrangement upon FcRn binding. (**E** – **G**) Schematic representation of the complex antibody formats with Format key in the three design principle groups. Format key: N: N-terminal Fab, C: C-terminal Fab, 0: no Fab. Fabs containing Binder#1 are indicated in green. (**H** – **J**) Bar graphs showing FcRn-recycling efficiency (%) as determined in the LSEC assay for antibody formats, grouped by design principles (**E** – **G**). Data are represented as mean +/- SD of n=3-15 independent experiments. Statistical analysis was performed using a one-way ANOVA [F(13, 66) = 51.74, *P <* 0.0001] followed by Tukey’s multiple comparisons test. ns = not significant, *: *P <* 0.05, **: *P* < 0.01, ***: *P* < 0.001. Data, exact n and *P* values are provided in Suppl. Table 1. (**K**) Violin plot showing the relative retention time (RRT) on a FcRn-affinity column indicating that the cell free environment does not well capture antibody format-dependent FcRn-recycling. Green diamonds: Binder#1 antibody formats, grey triangles: Canonical IgGs with Fv from marketed antibodies (RRT data from (21)). (**L**) Correlation of FcRn-recycling efficiency (from the LSEC assay) with the relative retention time (RRT) on a FcRn affinity column of complex antibody formats with Binder#1. Linear regression +/- confidence interval. Pearson correlation between cellular FcRn-recycling efficiency and FcRn relative retention time (r = 0.93, P < 0.0001, n = 13). Colored circles indicate the number of Fabs contained in a complex antibody format. Data and statistical analyses are provided in Table S1.

To evaluate these formats in a physiologically relevant environment, we employed a newly established (primary human) LSEC assay (Figure S1C), which preserves the endosomal membrane topology required to observe architectural constraints. In this assay, antibodies were labeled site-specifically via their Fc-glycosylation with pH-sensitive fluorophores and incubated with LSECs leading to non-specific endocytosis, and either rescue by FcRn-recycling or lysosomal degradation, causing fluorophore accumulation and fluorescence increase upon a drop in pH. By comparing the lysosomal accumulation of this fluorophore between two conditions, pharmacological block of FcRn (no FcRn-recycling) and a non-blocked condition (active FcRn-recycling), via flow cytometry, we quantified the recycled fraction of each antibody and related it to the total amount of antibody taken up, further called FcRn-recycling efficiency.

Consistent with our hypothesis, increasing the number of Fab domains at the N-terminus resulted in a stepwise decrease in FcRn-recycling efficiency. Compared with the parental, canonical IgG format (N-N), which exhibited FcRn-recycling efficiency of ∼33%, higher valency N-terminal formats showed a reduction to a minimum of ∼11% in the NN-NN format (approx. N-N > N-NN > N-NNN = NN-NN) (Figure 1H). In contrast, rearranging Fabs to the C-terminus (‘invertbodies’) significantly enhanced recycling efficiency, reaching ∼67% for the C-C format (approx. C-C > C-CC > CC-CC > N-N > N-NN > NN-NN) (Figure 1I). While C-CC and CC-CC variants demonstrated markedly improved recycling compared with their N-terminal counterparts (N-NN and NN-NN), a stepwise decrease in recycling efficiency relative to the C-C format was observed, suggesting that extreme bulkiness may still exert an influence even at the C-terminus. Finally, one-armed molecules also showed a significant increase in FcRn-recycling efficiency compared to canonical IgG (approx. 0-N > 0-NN = 0-NNN > N-N) (Figure 1J), suggesting that one-armed arrangement at the N-terminus also alleviates steric constraints. In line with the steric hindrance model, the contorsbody, with a distorted arrangement of the N-N arms, and the 0-NC format, with one N- and one C-terminal Fab, also led to an increased FcRn-recycling compared to the canonical IgG, while the mixed N- and C-terminal formats show similar (N-NC) or reduced (NC-NC) FcRn-recycling (Figure S1D).

To confirm that these differences were specifically driven by the presence of a membrane, we tested the antibody formats using cell-free FcRn-affinity chromatography (40) and, as an additional confirmation for the importance of the membrane constraints, an MDCK-based pulse-chase assay with overexpression of human FcRn (41). As anticipated for a system lacking a cell membrane, the FcRn-affinity column showed only minor differences in relative retention (RRT) (Figure 1K), located in a narrow range when compared to canonical IgG1 formats with Fv-sequences of marketed antibodies (21). However, these minor differences correlated strongly with the corresponding FcRn-recycling efficiencies of the LSEC assay (Figure 1L), indicating that the FcRn-column shows a small degree of steric hindrance, potentially caused by the bead surface to which FcRn is immobilized, though with minor effect compared to other factors mediated by the Fv-sequence. The pulse-chase assay on the other hand, which involves both FcRn-dependent uptake (pulse at lower pH) and sorting (endosomal pH) followed by extracellular release at neutral pH, qualitatively confirmed that the antibody format impacts FcRn-recycling efficiency in a similar manner for all three design rules (Figure S2A–C).

Together, these results demonstrate that the antibody architecture is a primary, quantifiable determinant of FcRn-recycling.

### 2. Antibody architecture-dependent FcRn-recycling is generalizable across variable domains

To ensure that the identified architectural rules are a fundamental property of the antibody format and not specific to a variable domain sequence, we assessed the generalizability of our observations using an unrelated Fv-sequence.

To test this, we repeated the in vitro FcRn-recycling efficiency assessment with the LSEC assay using a second panel of complex antibody formats comprising an unrelated variable domain (Binder#2), again without a target in humans or animals (Figure S3). This Binder#2 panel included two additional formats (0-C and C-NC), while lacking the contorsbody (Figure S3C).

The results from the Binder#2 antibody panel supported the format-dependent FcRn-recycling relationship established for Binder#1 antibodies, confirming our core design findings: I) Increased N-terminal size leads to reduced recycling, II) C-terminal Fab location increases FcRn-recycling III) as well as the one-armed antibodies (Figure S5A–D).

To extract the generalization of the format effect, we compared FcRn-recycling efficiencies of the 13 shared formats between both binders. Binder#1 and Binder#2 antibodies showed a very strong positive correlation (Pearson’s R = 0.96) in FcRn-recycling efficiency, highlighting the consistency between format-dependent recycling despite the unrelated Fv-sequences (Figure 2A). Additional statistical analysis via a two-way ANOVA, delineating the relative contributions of the Fv-sequence versus antibody format, further revealed that the format accounted for 84.5% of the total variance in FcRn-recycling efficiency (P < 0.0001), whereas the contribution of the specific binder sequence was not significant (0.06%, P = 0.51) (Figure 2B and C).

**Figure 2.**
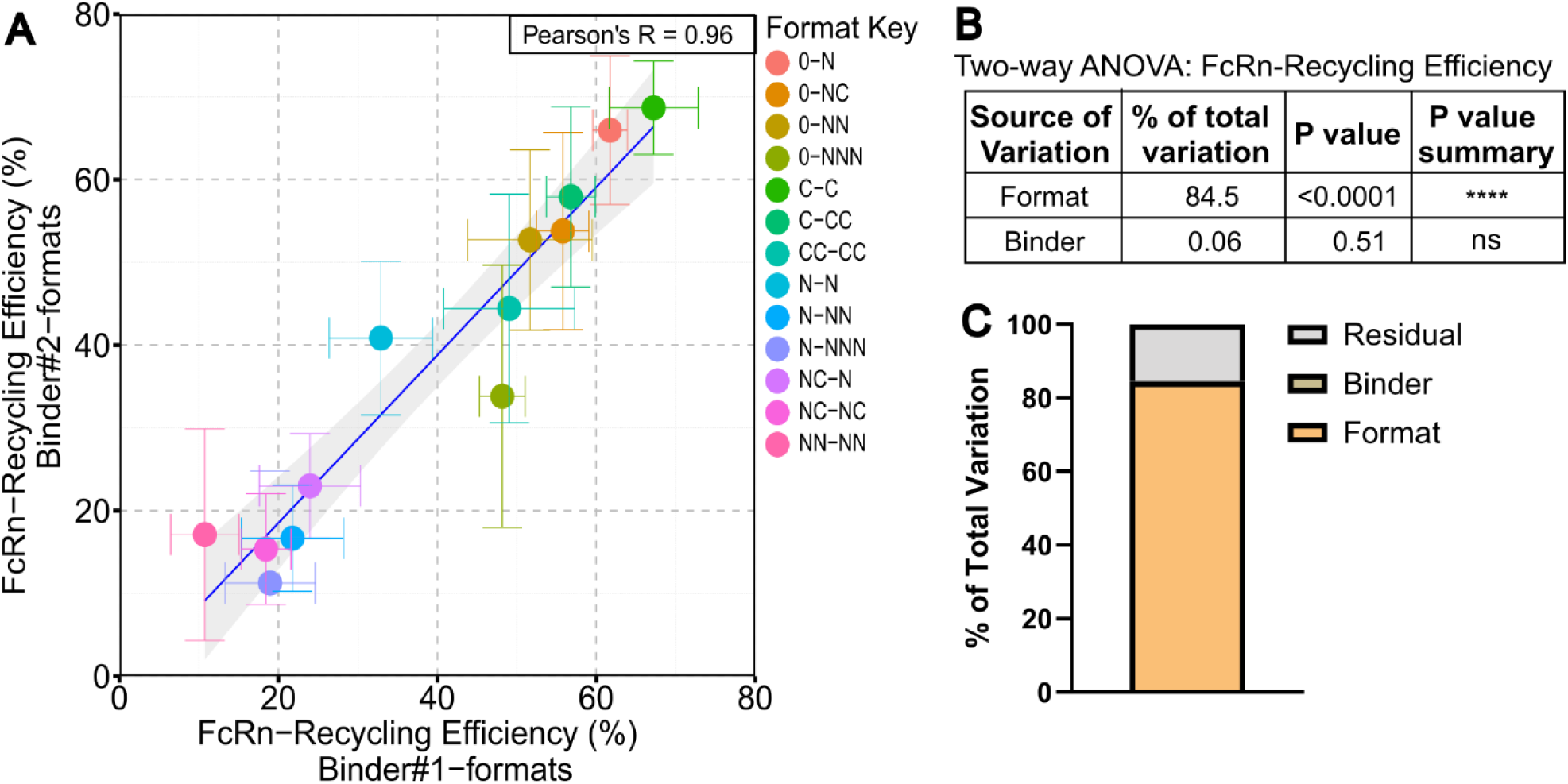
The antibody architecture, not the Fv-sequence, is the primary driver of FcRn-recycling efficiency. (**A**) Plot showing the correlation of FcRn-recycling efficiencies between matching formats built with Binder#1 and Binder#2. Pearson correlation analysis (R = 0.96, *P <* 0.0001, n = 13). Two-sided test. All data are mean +/- SD from n = 3-4 (Binder#2) or n = 3-15 (Binder#1) experiments. Linear regression (blue line) with 95% confidence interval (grey area) is shown. (**B** and **C**) Results of a statistical analysis using a two-way ANOVA to evaluate the effects of Format and Binder on FcRn-recycling efficiency. The analysis revealed a significant effect of the Format [*F*(*12, 113*) *= 51.46, P < 0.001*], but no significant effect of the Binder [*F*(*1, 113*) *= 0.44*, *P = 0.51*]. Data and statistical information are provided in Table S1.

These findings demonstrate that the architecture-intrinsic FcRn-recycling efficiency is a parameter that is translatable across antibodies with unrelated Fv-sequences, confirming that architectural design is a primary determinant of FcRn-recycling potential.

### 3. Fv-sequence and Fab number, not architecture, drive non-specific uptake

To fully deconstruct the biological processes contributing to non-specific clearance, we investigated whether initial cellular entry is similarly dictated by the antibody format or if it is governed by the biophysical properties of the variable domains (Fv). Since cellular uptake, the combination of fluid-phase and adsorptive endocytosis, is with the latter process driven by the interaction between the antibody’s surface and the cell membrane (18, 19, 21, 22, 42), we hypothesized that, unlike the architecture-dependency observed for FcRn-recycling, non-specific cellular uptake remains predominantly dependent on the specific Fv-sequence. To mechanistically isolate the uptake process, we utilized the FcRn-blocked condition of the LSEC assay, where lysosomal accumulation reflects the total amount of endocytosed antibody (Figure S1C). We compared the relative uptake across the formats for both Binder#1 and Binder#2 and complemented these cellular findings with heparin affinity chromatography as a surrogate for charge-mediated adsorptive endocytosis (21).

In contrast to the consistency observed in FcRn-recycling efficiency, the two unrelated binders exerted distinct effects in how the format affected the relative uptake (Figure S4A–D, Figure S5E–H). Direct comparison of matching Binder#1 and Binder#2 formats was not significantly correlated (Pearson’s R = -0.46, P = 0.11) (Figure 3A). However, when analyzed individually, Binder#1 showed a stepwise increase in relative uptake that did not relate with the spatial architecture, but with the number of Fab domains (ranking: 1 Fab < 2 Fabs < 3 ∼ 4 Fabs). In contrast, Binder#2 formats displayed lower overall uptake with a trend toward decreased relative uptake with higher number of Fabs, which was most pronounced in formats containing four Fabs (Figure 3B). These binder-dependent profiles were mirrored by the heparin affinity chromatography (21) results; while Binder#2 exhibited similar, low retention times across all formats, Binder#1 showed a valency-dependent increase in retention (Figure 3C). Statistical analysis of the 13 formats present with both Fv-sequences via two-way ANOVA further confirmed that the Fv-sequence was the primary driver of uptake variance (62%), whereas the complex antibody format contributed only 6% (Figure 3D and E), likely originating from the number of Fabs, rather than the architecture.

**Figure 3.**
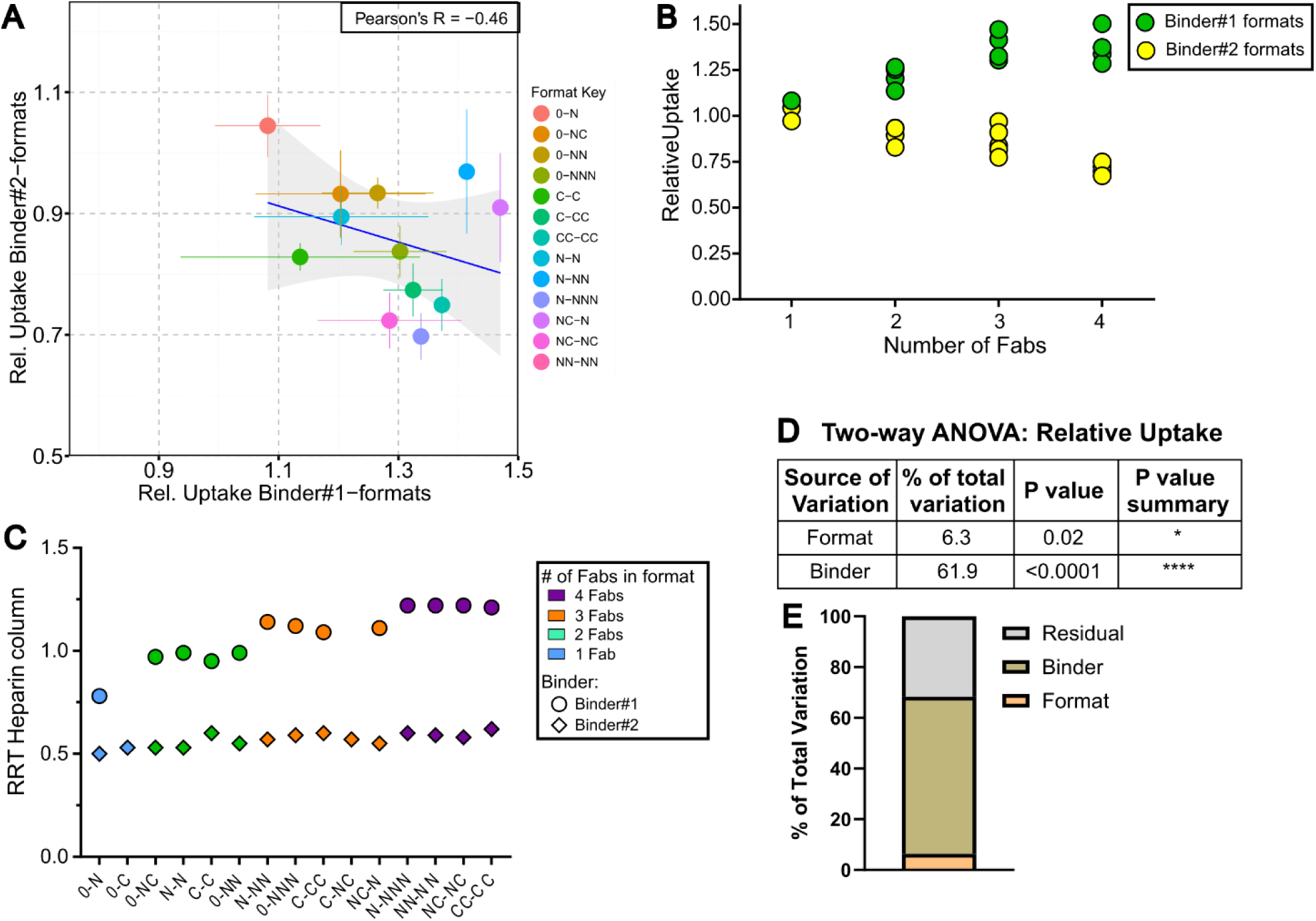
Fv-sequence and number of Fabs in a complex format affect non-specific endocytosis. (**A**) Plot showing the correlation of relative uptake between matching Binder#1 and #2 antibody formats. Pearson correlation analysis (R =- 0.46, *P = 0.11*, n = 13). Two-sided test. All data are mean +/- SD from n=3-4 (Binder#2) or n=3-15 (Binder#1) experiments. Linear regression (blue line) with 95% confidence interval (grey area) is shown. (**B**) Plot showing the relative uptake of complex formats, grouped by binder (color) and number of Fabs. (**C**) Relative retention time (RRT) on a Heparin affinity column for the different formats with Binder#1 and Binder#2. Colors represent different numbers of Fabs per format. (**D** and **E**) Results of a statistical analysis using a two-way ANOVA to evaluate the effects of Format and Binder on relative uptake. The results showed significant effects for both the Format [*F*(*12, 112*) *= 2.16*, *P = 0.018*] and the Binder [*F*(*1, 112*) *= 253.28*, *P < 0.0001*]. Data and statistical information are provided in Table S1.

Collectively, these results demonstrate that the complex antibody format influences non-specific endocytosis predominantly by the variable domain sequence and the resulting avidity-like effects of multiple Fab domains, rather than the spatial architecture *per se*. This emphasizes that architectural engineering specifically modulates the FcRn-recycling, with only limited impact on uptake.

### 4. Antibody format governs non-specific clearance in vitro and in vivo

While understanding the individual format effects on endocytosis and FcRn-recycling is crucial for optimizing the antibody, the translation of these mechanistic findings into clinically relevant insight requires understanding how both pathways converge on non-specific CL.

To capture this functional equilibrium in vitro, we utilized a third readout of the LSEC assay - relative cellular accumulation (rCA), measured under physiological conditions (non-FcRn blocked) to quantify the lysosomal antibody disposition (Figure S1C) - and quantified the rCA across all antibody formats for both Binder#1 and Binder#2. These experiments confirmed that the impact of the antibody format on rCA is fundamental, as both binder panels exhibited significant, format-dependent differences in rCA (Figure S6A–D). In particular, the Binder#1 formats demonstrated a clear pattern according to the design rules: I) increasing N-terminal size, elevates the rCA and thus the antibody elimination, while II) C-terminally located Fabs and III) one-armed antibodies reduced rCA when compared to the canonical IgG format. The Binder#2 formats followed the same overall trend (Figure S6E–H).

To dissect the individual impact of FcRn-recycling and relative uptake on rCA, we assessed each cellular mechanism against the corresponding rCA. For both Binder#1 and #2 binder panels, the FcRn-recycling efficiency correlated strongly with the rCA (Binder#1: Pearson’s R = -0.98; Binder#2: Pearson’s R = -0.89; (Figure 4A)), indicating that in both cases FcRn-recycling dictates differences in rCA. In contrast, the relative uptake data (Figure 4B) for Binder#1 antibodies correlated with a systematic increase in rCA (R = 0.76), whereas Binder#2 showed no such correlation (R = -0.11), highlighting that Fv-mediated uptake influences primarily the magnitude of cellular accumulation rather than the format-dependent hierarchy (Figure 4B). Most crucially, the rCA values for Binder#1 and #2 showed a strong linear correlation (Pearson’s R = 0.80; Figure 4C), establishing that the format-driven hierarchy remains constant across unrelated Fv-sequences.

**Figure 4.**
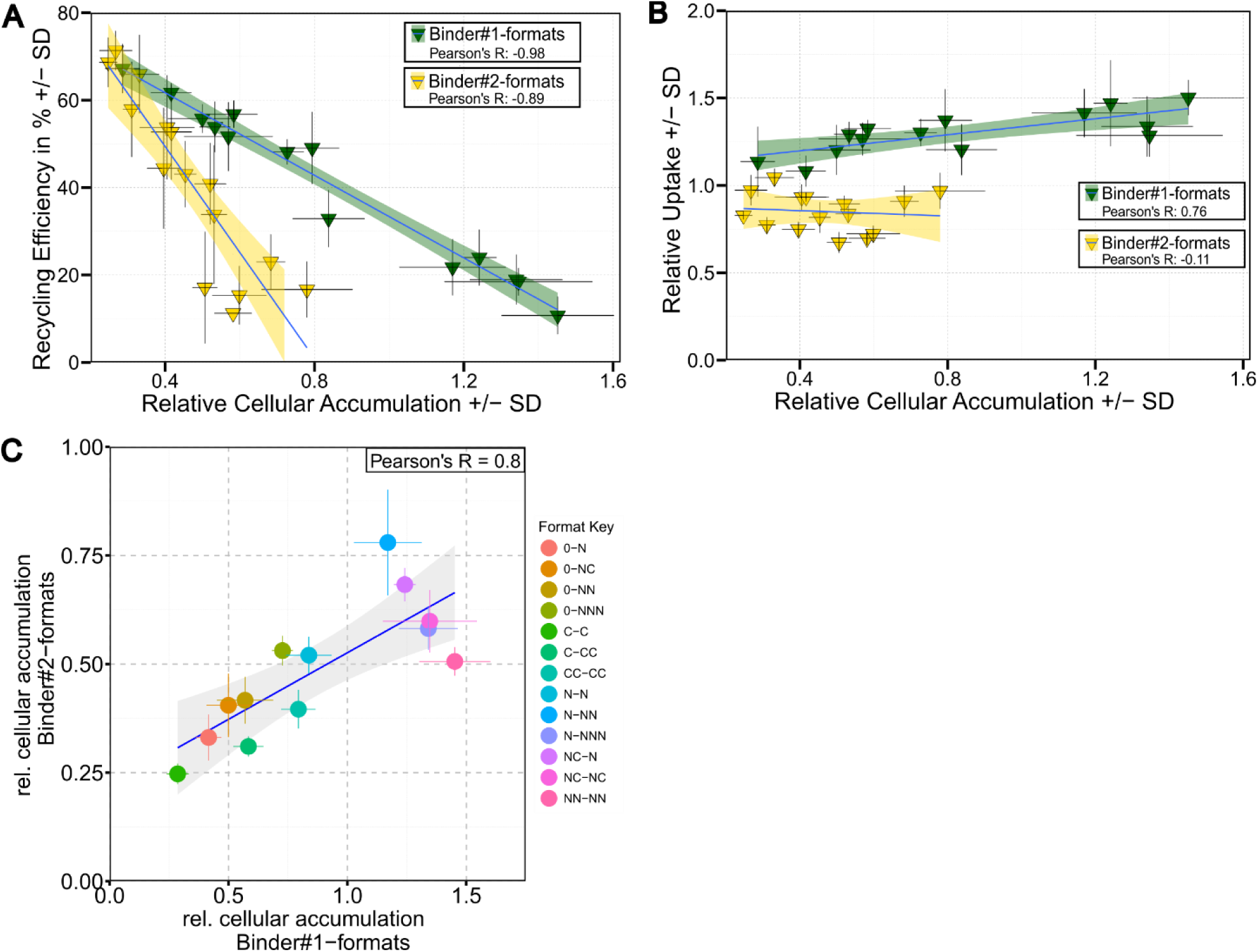
The antibody format dictates rCA differences primarily via FcRn-recycling. (**A**) Plot depicting the correlation between in vitro FcRn-recycling efficiency and the relative cellular accumulation (rCA) for formats with Binder#1 (green) and Binder#2 (yellow). Pearson correlation analysis of FcRn recycling efficiency versus rCA for Binder#1 (R = -0.98, *P <* 0.0001, n = 14) and Binder#2 (R = -0.89, *P <* 0.001, n = 15). Analysis was performed using two-sided tests. (**B**) Plot showing the correlation between the relative uptake and the rCA of formats with Binder#1 and Binder#2. Each triangle corresponds to an individual antibody format. Pearson correlation between relative uptake and rCA for Binder#1 (R = 0.76, P = 0.0018, n = 14) and Binder#2 (R = -0.11, P = 0.71, n = 15). Shaded areas represent the 95% confidence intervals of the linear regressions. (**C**) Plot showing the correlation of rCA between matching Binder#1 and #2 antibody formats. Pearson correlation analysis (R =0.8, *P < 0.001*, n = 13). Two-sided test. All data are mean +/- SD from n=3-4 (Binder#2) or n=3-19 (Binder#1) experiments. Linear regression (blue line) with 95% confidence interval (grey area) is shown. Data and statistical information are provided in Table S1

Building on these results, we validated the in vitro findings for eight representative (and two additional) Binder#1 formats across the design rules in vivo by performing single-dose pharmacokinetic (SDPK) studies in hFcRn Tg32 mice with coadministration of IVIG (Figure 5A, Table 1). These studies demonstrated that the translatability of the format driven antibody disposition can be recapitulated in vivo for the dual-armed Binder#1 antibodies. We observed a dramatic increase in clearance with increasing N-terminal number of Fab domains (N-N < N-NN < NN-NN, (Figure 5B and C)), while C-terminal location of Fabs (invertbody format) significantly reduced CL (C-C << N-N, C-CC << N-NN, CC-CC << NN-NN, (Figure 5D and E)). These in vivo results closely mirrored our in vitro rCA observations, confirming the format-specific impact on non-specific CL across the tested antibody architectures. The strong relationship was underscored by a robust linear correlation between the CL in hFcRn Tg32 mice and the in vitro assessed rCA of the dual-armed antibody formats (Pearson’s R = 0.92, Figure 5H).

**Figure 5.**
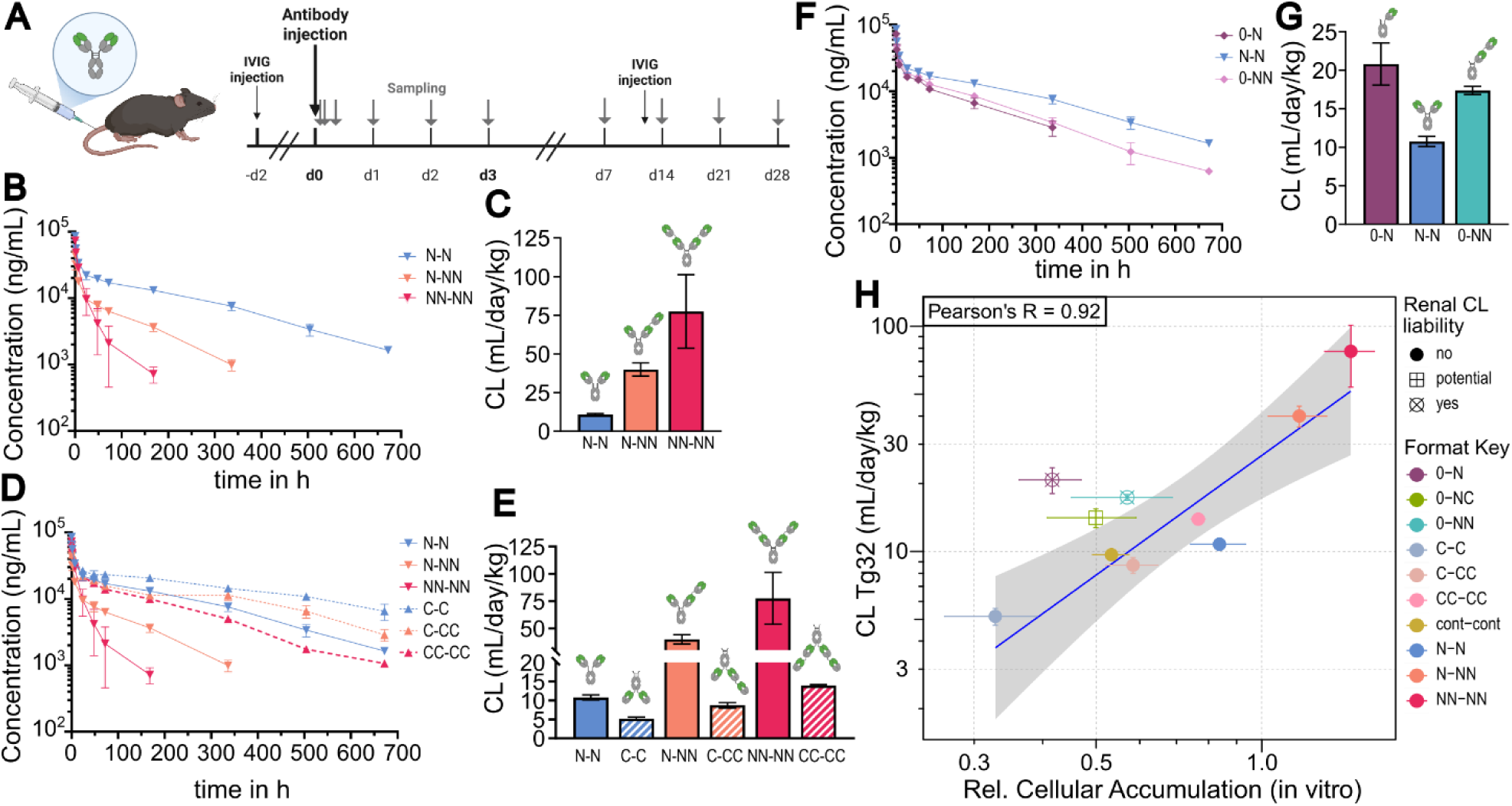
Mouse SDPK studies demonstrate strong influences of the antibody format on non-specific CL. (**A**) Schematic representation of the SDPK study design in hFcRn Tg32 mice. (**B**, **D** and **F**) Serum concentration-time profiles and (**C**, **E** and **G**) clearance (CL) in hFcRn Tg32 mice with IVIG coadministration for Binder#1 antibody formats grouped by the design principles from Figure 1. (**B** and **C**) increasing N-terminal size, (**D** and **E**) C-terminal Fab location, (**F** and **G**) one-armed antibodies. Data are shown as mean +/- SD for n=3-4 mice. (**H**) Plot depicting the correlation of the CL in mice with the rCA of each antibody format. One-armed antibody formats with renal CL liability were excluded from the regression and correlation analysis. Blue line and grey area show the linear regression with the 95% confidence interval. Pearson correlation analysis between relative cellular accumulation (Binder#1) and observed clearance in hFcRn Tg32 mice (R = 0.92, P = 0.0032, n = 7).

**Table 1:** Clearance of Binder#1 antibody formats in hFcRn Tg32 mice with IVIG co-administration and in FcRn-KO (b2m-KO) mice. n.t. - not tested

|  | huFcRn Tg32 +IVIG<br>CL (mL/day/kg) |  |  |  | FcRn-KO (b2m-KO)<br>CL (mL/day/kg) |  |  |  |
|---|---|---|---|---|---|---|---|---|
| **Format Key** | N | Mean | SD | CV% | N | Mean | SD | CV% |
| N-NN | 4 | 40.0 | 4.3 | 10.7 | 4 | 113.8 | 22.9 | 20.1 |
| 0-N | 4 | 20.8 | 2.7 | 13.1 | 4 | 89.6 | 10.2 | 11.3 |
| 0-NN | 4 | 17.4 | 0.5 | 3 | 4 | 100.8 | 3.3 | 3.2 |
| N-N | 3 | 10.8 | 0.7 | 6.2 | 4 | 84.0 | 4.1 | 4.9 |
| 0-NC | 4 | 14.1 | 1.3 | 9.5 | n.t | n.t. | n.t. | n.t. |
| NN-NN | 4 | 77.5 | 23.8 | 30.7 | 4 | 124.5 | 12.9 | 10.4 |
| C-C | 4 | 5.2 | 0.4 | 8.5 | 4 | 77.2 | 10.3 | 13.3 |
| cont-cont | 4 | 9.7 | 0.5 | 5.6 | n.t. | n.t. | n.t. | n.t. |
| C-CC | 4 | 8.7 | 0.7 | 8 | 4 | 76.2 | 11.4 | 15.0 |
| CC-CC | 3 | 13.9 | 0.3 | 1.8 | 4 | 104.5 | 5.6 | 5.3 |

In contrast, one-armed architectures (0-N and 0-NN) – designed to alleviate steric hindrance upon FcRn engagement through an expanded accessible conformational space, and which exhibited low rCA in vitro in line with increased FcRn-recycling efficiencies – had unexpectedly high CL compared to the canonical IgG format (N-N) (Figure 5F–G). This discrepancy between in vitro rCA and in vivo outcome likely stems from the fact that exclusively one-armed antibody formats are susceptible to glomerular filtration – a format-specific renal clearance mechanism that is not captured by our in vitro LSEC assays but contributes significantly to the total clearance of one-armed antibody formats in different preclinical species and in humans (43–46).

Finally, we performed a mechanistic decomposition of the two non-specific CL processes in vivo. To this end, we conducted SDPK studies in FcRn knock-out (FcRn-KO) mice (Figure 6A), which lack FcRn-recycling due to the deletion of the ꞵ2-microglobulin encoding gene. In these animals, non-specific CL is primarily driven by cellular uptake. While all assessed antibody formats exhibited the high clearance characteristic of FcRn-KO mice (Figure 6B and C, Table 1) the variation between the individual formats was narrow. Specifically, the CL varied by a maximum of only 1.6-fold - ranging from 76.2 to 124.5 mL/d/kg - which correlated well with the in vitro assessed relative uptake of dual-armed antibodies (Pearson’s R = 0.85, Figure S7). In contrast, the CL range in hFcRn Tg32 mice, where FcRn-recycling is active, spanned from 5.2 to 77.5 mL/d/kg, representing a 15-fold difference (Figure 6D). Comparing these datasets (15-fold vs. 1.6-fold) highlights an approx. 10-fold window of CL reduction mediated directly by FcRn. To corroborate this mechanism, we calculated an in vivo recycling efficiency (CL_FcRn-KO_ / CL_hFcRn Tg32_), which correlated strongly with the in vitro FcRn-recycling efficiency for the dual-armed antibodies (Pearson’s R = 0.93; Figure 6E). This mechanistic deconvolution confirms the antibody format FcRn-recycling relationship in vivo and highlights the predictivity of the in vitro data.

**Figure 6.**
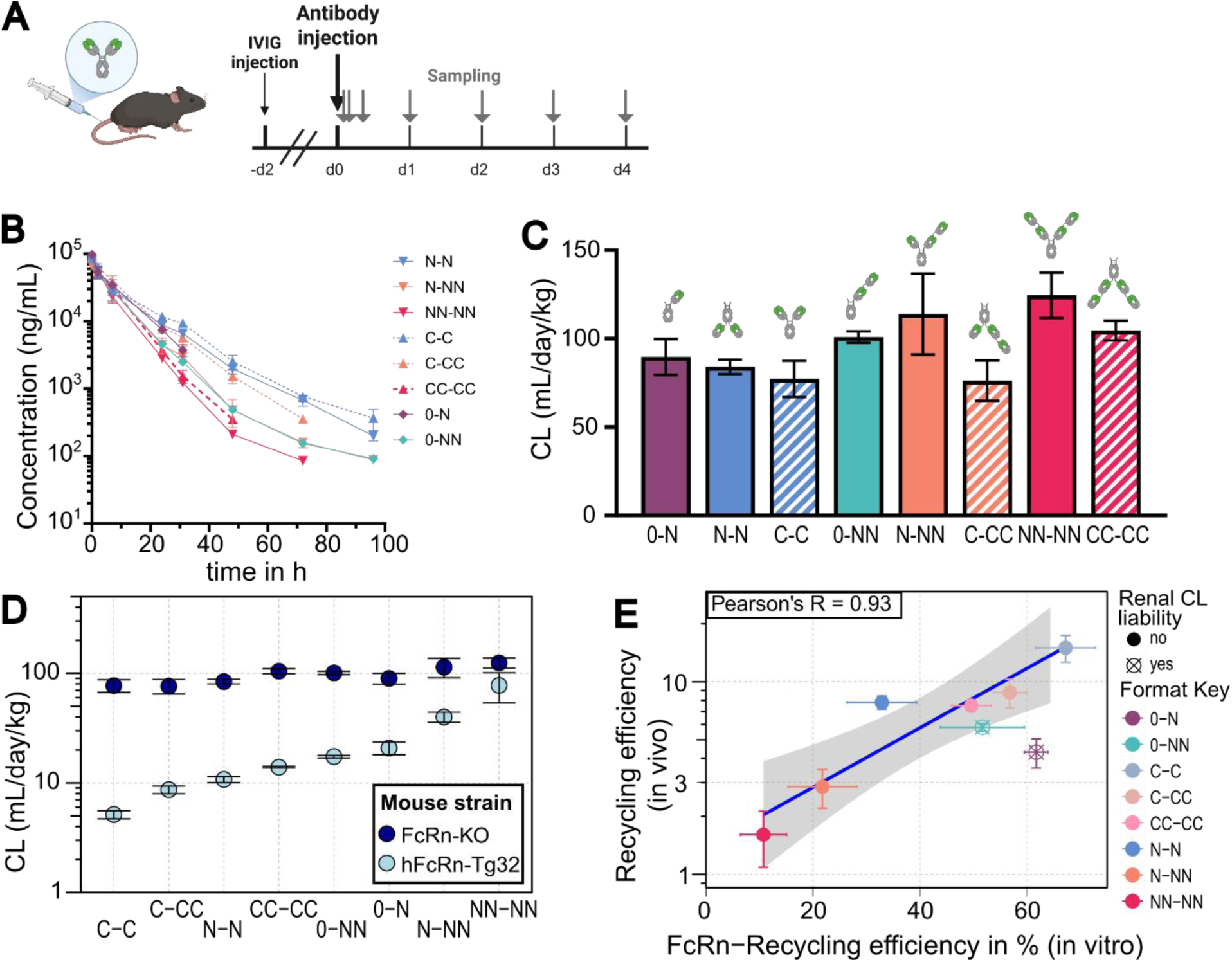
In vivo mechanistic decomposition confirms the antibody architecture-dependent FcRn-recycling. (**A**) Schematic representation of the SDPK study in FcRn-KO (b2m-KO) mice. (**B**) Serum concentration-time profiles of different Binder#1 antibody formats in FcRn-KO mice. Data are presented as mean +/- SD for each time point. (**C**) Corresponding CL of the Binder#1 antibody formats in FcRn-KO mice. n = 4. (**D**) Graph comparing the CL in hFcRn Tg32 and FcRn-KO mice, arranged by format. Mean +/- SD. (**E**) Plot depicting the correlation between in vitro and in vivo FcRn-recycling efficiencies. Pearson correlation between in vitro and in vivo FcRn-recycling efficiencies (R = 0.93, P = 0.0076, n = 6). One-armed antibody formats with a liability for renal catabolism are indicated and excluded from regression and correlation analyses. Blue line + grey area shows the linear regression with the 95% confidence interval. Pearson correlation R is indicated in the top left. Two-sided tests were used.

Collectively, these findings establish that the antibody architecture is a predictable determinant of non-specific CL by driving FcRn-recycling, while demonstrating robust translatability of the in vitro findings to in vivo for dual-armed antibodies.

### 5. Architectural engineering complements Fc-FcRn-mediated affinity optimization to maximize FcRn-recycling efficiency

Having demonstrated that the antibody architecture heavily influences FcRn-recycling and consequently non-specific CL, we next evaluated whether this structural engineering can be combined with affinity-based engineering. Traditional half-life extension strategies use Fc-mutations to increase binding affinity to FcRn to increase recycling, such as the ACT5 variant (M428L/N434A/Q438R/S440E) (28, 29). Hypothesizing that the antibody format and Fc-FcRn engineering act as independent, additive variables, we generated six Binder#1 antibody formats, incorporating the ACT5 mutations. We evaluated these engineered variants in comparison to the wt-variants to determine if affinity-based optimization could override the steric liabilities of complex antibody architectures or whether combining both strategies can yield a synergistic ‘super-recycling’ phenotype.

We utilized the LSEC assay to measure the in vitro FcRn-recycling efficiency of the six Binder#1 variants (N-N, N-NN, NN-NN, C-C, 0-N, 0-NN) in both ‘Fc-WT’ (without an Fc-FcRn engineering) and ACT5-engineered forms. Incorporating the ACT5 mutations resulted in an increased FcRn-recycling efficiency for every tested format compared to its Fc-WT (without affinity engineering) counterpart (Figure 7A). However, the relative differences between ACT5 engineered and Fc-WT formats were reduced in a predictable format-dependent manner (NN-NN > N-NN > N-N > 0-NN > 0-N > C-C). Formats with high FcRn-recycling efficiencies benefited less from the additional Fc-FcRn engineering mutations.

**Figure 7.**
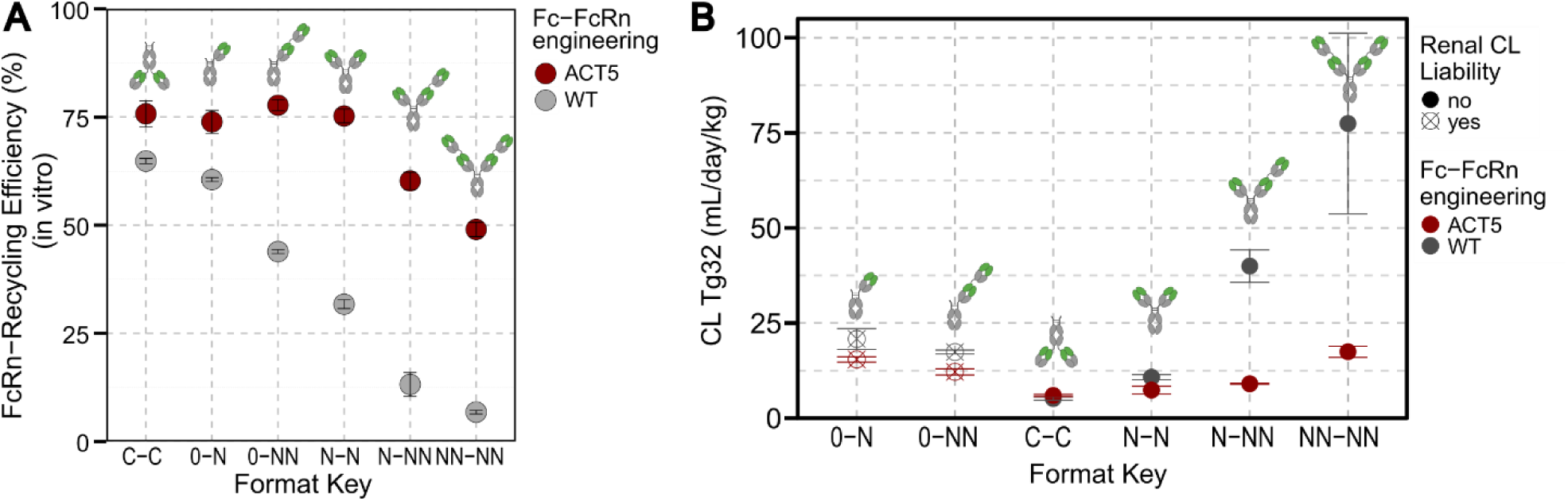
The impact of Fc-FcRn engineering depends on the antibody architecture. (**A**) FcRn-recycling efficiencies (in vitro) of Binder#1 antibody formats with Fc-FcRn engineering (‘Fc-ACT5’, red) and without (‘Fc-WT’, grey). Data represent the mean +/- SD from n=3 independent experiments. (**B**) Serum clearance determined in hFcRn Tg32 mice with IVIG coadministration for antibody formats without Fc-FcRn engineering (grey, from Figure 5) and with ACT5 mutations (red). One-armed antibody formats with a liability for renal catabolism after glomerular filtration are indicated. Data are mean +/- SD (n=3-4 mice per group). Data are provided in Table S1

Additionally, while the architectural hierarchy for FcRn-recycling was preserved in the presence of the ACT5 mutation for the N-terminal NN-NN (∼49%), N-NN (∼60%) and the canonical IgG N-N format (∼75%), the invertbody (C-C) and one-armed antibody formats (0-N, 0-NN) did not exhibit further increases in FcRn-recycling efficiency compared to the canonical format, indicating that the assay reached a ‘recycling ceiling’ for these high-recycling antibodies. These in vitro analyses clearly demonstrate that architectural engineering and FcRn-affinity optimization act as complementary strategies with an upper limit.

To evaluate if this ‘recycling ceiling’ translates to non-specific CL in vivo, we performed SDPK studies with the ACT5-engineered antibodies in hFcRn Tg32 mice. The observed CL values (Table 2) recapitulated the in vitro FcRn-recycling efficiencies (Figure 7B) for dual-armed antibodies, confirming that FcRn-affinity optimization significantly reduces non-specific CL, whereas the absolute limit is governed by the antibody architecture. For low FcRn-recycling formats such as the NN-NN, the ACT5 mutations reduced clearance by over four-fold (17.5 mL/d/kg vs. 77.5 mL/d/kg for Fc-WT), yet it remained still higher than the canonical IgG without FcRn-engineering (∼10.8 mL/d/kg) (Figure 7B). Conversely, the C-C invertbody format, which already possesses an optimized spatial orientation, showed no further CL reduction with the addition of ACT5 (6.0 mL/d/kg vs. 5.2 mL/d/kg for Fc-WT), essentially reaching the “recycling ceiling” in the in vivo model (Figure 7B). The one-armed variants (0-N, 0-NN) showed intermediate CL reduction by ACT5 mutations, but again had higher CL than the canonical IgG (N-N) without ACT5 mutations, presumably reflecting the presence of an additional renal clearance pathway (43).

**Table 2:** Clearance of Binder#1 antibody formats with ACT5 Fc-FcRn engineering in hFcRn Tg32 mice with IVIG co-administration.

|  | huFcRn Tg32 +IVIG<br>CL (mL/day/kg) |  |  |  |
| **Format Key** | N | Mean | SD | CV% |
|---|---|---|---|---|
| N-N (ACT5) | 4 | 7.4 | 1.0 | 14.1 |
| N-NN (ACT5) | 4 | 9.0 | 0.2 | 2.0 |
| 0-N (ACT5) | 4 | 15.4 | 0.7 | 4.5 |
| 0-NN (ACT5) | 4 | 12.2 | 0.8 | 6.8 |
| NN-NN (ACT5) | 3 | 17.5 | 1.5 | 8.5 |
| C-C (ACT5) | 4 | 6.0 | 0.3 | 4.9 |

In summary, these findings establish that optimizing non-specific CL of therapeutic antibodies via FcRn-recycling efficiency is most effective using a dual strategy: architectural engineering to alleviate fundamental steric constraints and Fc-FcRn engineering to maximize the efficiency of neonatal receptor binding.

### 6. An invertbody variant of Golimumab improves pharmacokinetics without compromising activity

To demonstrate that architectural engineering represents a viable and effective strategy for optimizing the non-specific clearance of therapeutic antibodies, we applied our design rules to the marketed anti-TNF*a* antibody golimumab (47, 48). We hypothesized that structurally inverting golimumab into a C-terminal invertbody (C-C) would alleviate endosomal membrane constraints and enhance FcRn-mediated recycling, potentially rivaling the performance of affinity-mediated Fc-FcRn engineering. To test this, we compared the parental golimumab to an inverted variant with a WT-Fc and an ACT5-engineered variant of the parental mAb. We first ensured that the structural modification did not impair functional potency. In vitro activity assays confirmed that moving the Fabs to the C-terminus did not compromise biological activity, as the inverted variant exhibited an equivalent potency in inhibiting TNF*a* signaling (IC50: 28 pM) compared to both the parental antibody (IC50: 38 pM) and the ACT5-variant (IC50: 31 pM) (Figure 8A). The subsequent evaluation of these variants in the LSEC assay confirmed that C-terminal Fab arrangement significantly enhances FcRn-recycling efficiency in a physiologically relevant environment. The inverted variant (70% recycling) markedly outperformed the parental golimumab (50% recycling) and reached a level similar to the ACT5 affinity-engineered control (71% recycling) (Figure 8B). These improvements in FcRn-recycling, assessed in vitro, translate directly into a substantial reduction in systemic clearance in hFcRn Tg32 mice with IVIG coadministration. The inverted variant exhibited a clearance of 8.9 mL/d/kg, representing a three-fold reduction compared to the parental antibody (27.6 mL/d/kg) and matching the performance of the ACT5-variant (10.4 mL/d/kg) (Figure 8C and D).

**Figure 8.**
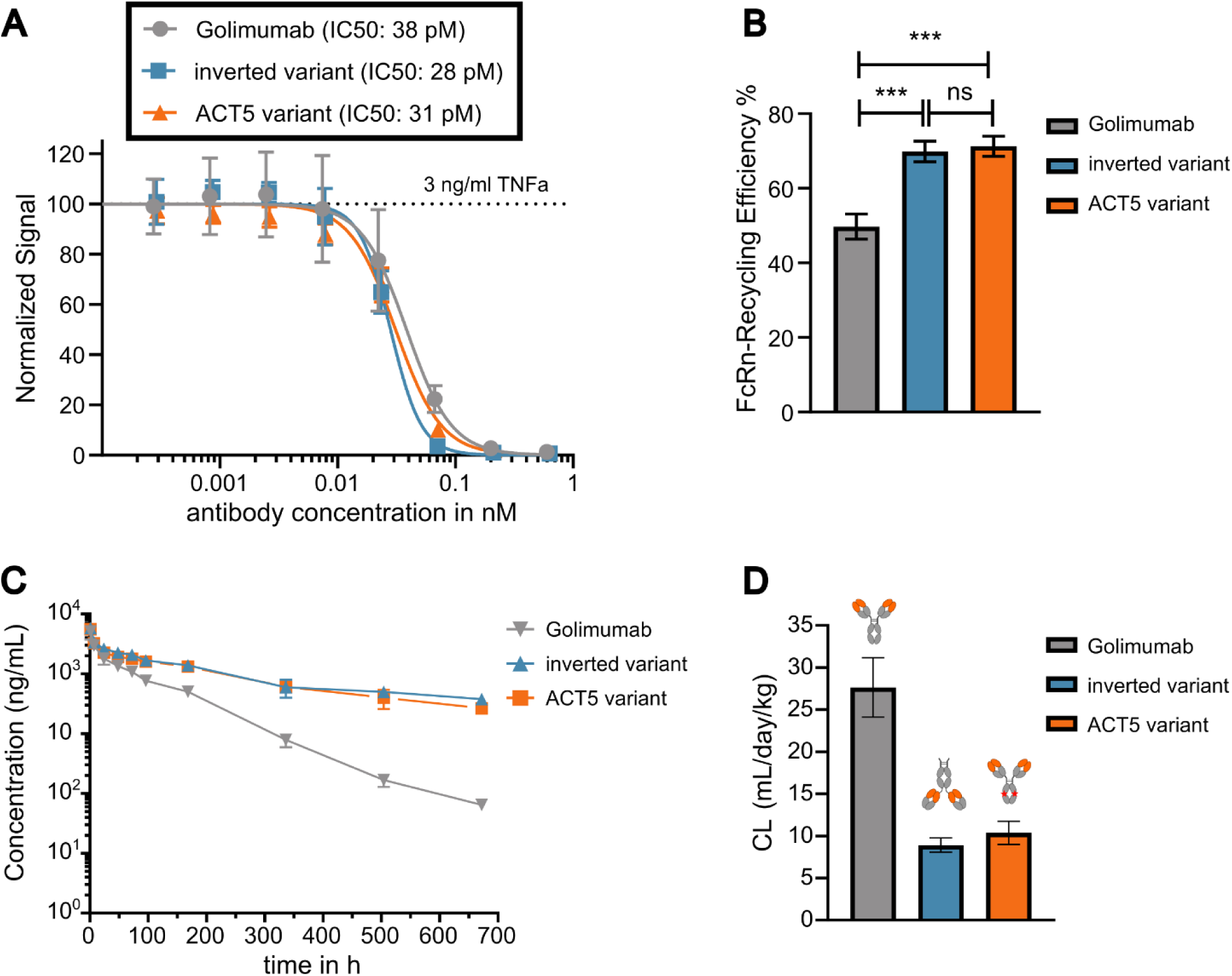
The inverted variant of a therapeutic anti-TNF-α antibody retains its inhibitory activity while showing a reduced non-specific clearance. (A) TNF-α neutralization by antibody variants. Dose-response curves for Golimumab, the inverted variant (C-C), and the ACT5 variant against 3 ng/mL TNF-α. The Y-axis shows the signal normalized to the TNF-α only control. Data are mean +/- SD from n=3 technical repeats. (B) FcRn-recycling efficiencies of Golimumab, the inverted variant, and the ACT5 variant. Data represent mean +/- SD (n=3). Statistical analysis by one-way ANOVA [F(2, 6) = 49.83, P = 0.0002] with Tukey’s multiple comparisons test. ns = not significant, ***: *P* < 0.001. (C) Serum concentration-time profiles and (D) CL of golimumab, the inverted, and the ACT5 variant determined in hFcRn Tg32 SDPK studies with IVIG coadministration. Data are mean +/- SD (n=3-4 mice per group). Data and statistical information are provided in Table S1.

Collectively, these results provide a case study for the application of architectural engineering on an approved antibody. By demonstrating that structural inversion can achieve CL reduction comparable to established affinity-based methods without loss of potency, we validate the invertbody as a powerful format for the next generation of antibody therapeutics.

## DISCUSSION

This investigation systematically evaluated a diverse panel of non-canonical antibody formats to establish that the spatial architecture fundamentally governs up to 6-fold differences in FcRn-mediated recycling. This structure-recycling relationship followed systematic design rules and translated with high magnitude to non-specific clearance, driving up to 15-fold variance in vivo. As antibody-based therapeutics accounted for 20-30% of all approved drugs in recent years (49), with currently over 100 diverse complex formats in clinical development (4), the established structure-PK relationship sheds light on an underappreciated feature of complex antibody formats. While decades of FcRn research have centered on Fc-affinity and pH-dependent binding kinetics as the principal determinants of IgG recycling, our systematic decoupling of architecture-driven FcRn-recycling from Fv-sequence-driven cellular uptake identifies antibody architecture as an independent axis of control over receptor-mediated salvage.

The core of our findings highlights that while complex antibodies are designed to address unmet therapeutic needs, e.g. through multi-valency or novel modes of action (1–3), their spatial architecture profoundly impacts baseline pharmacokinetics. We identified three architectural rules regarding PK properties:

1. Increased N-terminal size elevates non-specific clearance
2. Inverted, C-terminal Fab arrangement minimizes non-specific clearance
3. One-armed configurations maximize FcRn-recycling but incur additional renal CL.

These rules provide a rational basis for selecting or avoiding specific antibody architectures, allowing pharmacokinetic behavior to be anticipated from structural design rather than discovered empirically later in development.

Reframing the rationale that non-specific clearance is governed by biophysical properties of the variable domains (Fv) and Fc-FcRn affinity, our findings establish that the architecture acts as an additional strong and independent modulator dictating endosomal FcRn-recycling. Prior work has proposed a simple steric hindrance mechanism by which Fab domains in canonical IgG impede FcRn engagement at the endosomal membrane (25, 30, 31), but direct evidence for how strongly this mechanism governs FcRn-recycling efficiency or, consequently, the systemic clearance of an antibody itself – and whether its impact scales with Fab number and arrangement – remained unknown. We show that it is neither subtle nor incidental: it drives large, predictable, stepwise differences in both FcRn-recycling and non-specific clearance, matching the design rules illustrated in Figure 1A–D, and transforming a structurally plausible mechanism into an engineering-actionable determinant of antibody pharmacokinetics.

The systematic deconstruction of non-specific clearance allows us to propose a conceptual model:

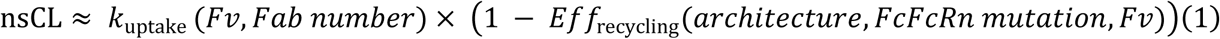

This framework separates non-specific CL (nsCL) into two independent processes, each with its own engineering levers: cellular uptake (k_uptake_), which is governed mainly by Fv biophysical properties and, secondarily, by Fab number; and FcRn-recycling efficiency (Eff_recycling_), governed primarily by antibody architecture and Fc-FcRn affinity engineering, but also by Fv properties.

Architecture and Fc-FcRn affinity engineering exert independent effects on FcRn-recycling (Eff_recycling_), yet their combined impact follows a clear pattern rather than simple additive effects. Testing the FcRn-affinity enhancing ACT5 mutations across the antibody format panel revealed a principle of diminishing returns: ACT5 markedly improved recycling in sterically hindered formats (N-NN and NN-NN), offered a moderate benefit in canonical IgG (N-N), and provided no further gain in the C-terminal invertbody (C-C). As a result, both invertbody variants (with and without ACT5) and ACT5-engineered canonical IgG converged on similar high recycling efficiency in vitro and the same minimal clearance in vivo, indicating a physiological ‘recycling ceiling’ beyond which neither strategy can further enhance FcRn salvage.

This recycling ceiling implies that even maximal endosomal FcRn salvage leaves a fraction of antibodies (∼25% in our assay) unrescued. Several non-exclusive mechanisms could contribute to this escape, including stochastic leakage of FcRn into the lysosomal degradation system (50), rapid endosomal transit upon gradual acidification, which compresses the window to reach maximum antibody-FcRn occupancy (51, 52), or limitations imposed by the pH dependent ‘affinity threshold’ restricting maximum FcRn-affinities: increasing pH 6 affinity beyond a certain point impairs pH 7.4 exocytic release and thus accelerates systemic clearance (53). Steric constraints in certain formats may lower this threshold, potentially allowing them to tolerate higher-affinity mutations lifting the format-dependent FcRn-recycling closer to the recycling ceiling – an open question for future studies.

Biophysical properties of the variable domains (Fv) contribute to both FcRn-recycling (Eff_recycling_) and non-specific endocytosis (k_uptake_). Positively charged Fv patches can stabilize the IgG-FcRn complex (54–56), while strongly negatively charged Fv domains have been linked to accelerated clearance in vivo (19), potentially via charge repulsion with the negatively charged endosomal membrane. We observed no such effects on FcRn-recycling between our two binder panels, a deliberate feature of our design: both Fv-sequences were chosen for typical, non-extreme biophysical properties to isolate properties mediated by the antibody architecture. For non-specific endocytosis (k_uptake_), by contrast, positively charged Fv patches drive adsorptive endocytosis via the negatively charged glycocalyx (18, 19, 21, 42, 57), an effect amplified by Fab number, since additional Fab domains multiply the number of charge patches available for membrane engagement, independent of their spatial arrangement. Fv charge optimization therefore becomes increasingly important as antibody valency increases.

Recapitulating the conceptual model with a theoretical example illustrates a further case of balancing Fv engineering against FcRn-recycling optimization: once FcRn-recycling is maximized – like in the invertbody, at the recycling ceiling – the rate-limiting step for non-specific clearance shifts entirely to cellular uptake. In practice this means that neither architecture nor affinity engineering can further improve an antibody which reached the recycling ceiling, but Fv engineering can. Non-specific CL optimization therefore cannot rely on a single lever through a molecule’s development – which lever matters depends on where on the Recycling-Uptake Axes the current antibody is located.

Applying our design rules to the marketed anti-TNF-α antibody golimumab confirmed the translational value. Re-engineering golimumab into a C-terminal invertbody variant reduced clearance in vivo – matching the benefit of the ACT5 mutation – without compromising in vitro potency. While binding potency was preserved in this case, it remains unclear whether C-terminal Fab positioning generally maintains affinity, or whether this is target- or epitope-dependent. This example demonstrates that architectural engineering is not merely a mechanistic finding but a viable tool for PK optimisation.

Two methodological choices were critical to uncovering this structure-PK relationship. First, membrane-competent cellular assays were essential: cell-free methods such as surface plasmon resonance (31) or FcRn-affinity chromatography (40) capture FcRn-affinity differences well, but the lack of the lipid-bilayer fails to capture the steric constraints, essential for comparison of different formats and thus critical for this study. Our LSEC assay, by contrast, achieved a strong in vitro-in vivo correlation (Pearson’s R = 0.92), underscoring the need for membrane-competent environments when evaluating non-specific CL properties of complex antibody formats, and offering a framework to reduce animal use during early PK de-risking, in line with regulatory 3R initiatives (58, 59). Second, a translational, physiologically competitive in vivo model was equally important: co-administering high-dose IVIG in hFcRn Tg32 mice exposes FcRn-recycling differences (6, 36) – whether driven by architecture or by affinity engineering – that would otherwise remain obscured by the mouse model’s naturally low endogenous IgG levels (35).

Finally, our SDPK studies revealed a clear in vitro in vivo mismatch for the one-armed antibody format (0-N and 0-NN), consistent with a distinct CL mechanism affecting exclusively one-armed antibodies: renal catabolism after glomerular filtration. Although antibody-sized proteins are traditionally considered too large to undergo renal elimination, Rafidi et al. demonstrated that shape and structural flexibility can enable glomerular passage for certain antibodies (43). Antibodies with a single Fab arm possess the conformational flexibility to adopt streamlined orientations that can pass through the glomerular pores, undergoing subsequent catabolism within proximal tubule cells (43). Consistent with this mechanism, studies across mice, non-human primates and humans demonstrate pronounced renal accumulation of radiolabeled one-armed antibodies (44–46). While the LSEC assay captures the superior FcRn-recycling of the one-armed designs, in vivo we can only observe total CL, the sum of different CL mechanisms, where in this case the additional renal clearance outweighs the gain from improved FcRn-mediated salvage. Notably, model-fitted clearance estimates from Rafidi et al. indicate a similarly elevated CL for analogous one-armed formats relative to bivalent IgG, in the order of 1.5- to 2-fold (43).

While this investigation establishes a new antibody intrinsic property relevant for target-independent, non-specific clearance, these findings must be contextualized within the multifactorial process of therapeutic antibody disposition. Saturable, molecule-specific pathways, such as target-mediated drug disposition (TMDD) or the development of anti-drug antibodies (ADA), can strongly accelerate systemic CL in ways that are inherently dependent on the biology of the target and therefore not generalizable (11).

Furthermore, while our structural panels focused exclusively on Fab-based arrangements around an IgG Fc scaffold, the biopharmaceutical landscape continues to diversify into alternative domains, including single-domain heavy-chain variables (VHH) (60). Although these alternative modalities fall outside the scope of this study, our core findings emphasize that any structural modification altering the spatial proximity to the endosomal membrane requires careful pharmacokinetic consideration and may serve as the basis for future investigations.

With more than 100 complex antibody formats now advancing through clinical development, our findings show that structural innovation comes with direct, predictable pharmacokinetic consequences driven by the spatial architecture. Defining how Fab positioning governs FcRn-recycling and non-specific clearance, we provide a generalizable framework that elevates antibody architecture from an afterthought to a controllable design variable. Embedding these principles into early antibody engineering should reduce PK-driven attrition and improve the translational success of next-generation biotherapeutics.

## MATERIALS AND METHODS

### 1. Antibody design and production

All complex antibody formats were designed using consistent antibody building blocks to isolate the effect of the architecture from other sequence-level variables. Across both Binder#1 and Binder#2 panels, all formats shared the same, standard Fc-domain sequence, the same GS-Linker for C-terminal or additional Fabs, and the same Fab sequences, with Fv-sequences for either Binder#1 or Binder#2. Except for the golimumab variants, all investigated antibodies carried the PGLALA mutation in the Fc-region to abolish effector functions. Where indicated, FcRn-affinity mutation ACT5 (M428L/N434A/Q438R/S440E) were introduced in the Fc-domains. All antibodies were produced in-house by transient transfection of Hek293 cells and purified using standard Protein A chromatography or were purchased from external partners (GeneArt or Proteros). Quality was assured by analytical size exclusion chromatography, CE-SDS, and/or mass-spectrometry analysis.

### 2. Antibody labelling for the LSEC assays

Antibodies were labeled using the SiteClick™ Antibody Azido Modification Kit (#S20026, Thermo Fisher Scientific) according to the manufacturer’s instructions. Briefly, N-linked galactose residues of the Fc region were removed by β-galactosidase and replaced by an azide-containing galactose (GalNaz) via the β-1,4-galactosyltransferase (GalT). The pH-sensitive amine-reactive dye pHAb (532 nm) was purchased from Promega (#G9841) and coupled to a sulfo DBCO PEG4 amine (synthesized internally). Antibodies were labeled with the DBCO modified pHAb by incubating the GalNaz modified antibodies with a molar dye excess of 3.5. Excess dye was removed using the Amicon® Ultra-2 Centrifugal Filter with a MWCO of 30 kDa (Merck, #UFC205024) and antibodies were re-buffered in 20 mM histidine 140 mM NaCl buffer (pH 5.5). The concentration of the labeled antibodies as well as the dye to antibody ratio were determined using a spectrometer (Lunatic Reader) via absorption measurement at 280 nm and 532 nm. Integrity of labelled antibodies and complete removal of excess dye was verified by analytical size exclusion chromatography (BioSuite Diol (OH) column, Waters, #186002165) in comparison to the parent, unlabeled antibody.

### 3. Assessment of relative cellular accumulation

Liver sinusoidal endothelial cells (LSECs) (HLECP2, LONZA, passage 4) were thawed and cultured at 37 °C and 5 % CO2 on collagen I coated dishes (Greiner, #354407) five days prior to the experiment in Endothelial Cell Growth Basal Medium-2 (Lonza, #CC-3156) supplemented with SingleQuots Supplements (Lonza, #CC-4176). Two days prior to treatment, cells were sub-cultured into collagen-I (Corning, #354407) coated 96-well plates at a cell density of 4×10×4 cells/well to allow adherence for 48 hours. On the day of the experiment, medium was replaced and cells were treated with a dosing solution of 200 nM labelled test antibodies in culture medium and incubated for 2 h and 4 h. The fluorescence of the dosing solution was measured in a Tecan Infinite Pro 300 fluorometer using 20 µl of dosing solution mixed with 60 µl citric acid buffer (0.2 M Citrate-Phosphate buffer pH 4.5) at an excitation wavelength of 532 nm and emission wavelength of 560 nm.

After incubation with the labelled antibodies, cells were washed twice with PBS and detached with 100 ul Accutase (PAN-Biotech, #P10-023100) for 2.5 min at 37°C. Detachment and antibody uptake processes were stopped with 100 ul of ice cold FACS buffer (20% FCS, 1 mM EDTA, PBS, 0.1 ug/ml DAPI) and cells were transferred into a fresh 96 well plate (Corning, #3596). Cellular accumulation of the labelled antibody was measured by FACS (BD LSR-Fortessa X20) using 532 nm excitation wavelength and recording emission at 560 nm. Raw FACS data were analyzed with FlowJo 10.10. Cells were gated for morphology, granularity, singlets, living (DAPI negative) and the geometric mean fluorescence intensity (MFI) was extracted from the final gate. The individual MFI for each sample at each time point was subtracted by the MFI of cells without antibody treatment. The background subtracted MFI were normalized to the fluorescence of the dosing solution to adjust for different dye to antibody ratios. The corrected MFI were used for a linear regression through origin (t = 0 h), the 2 h and the 4 h values. Slopes were used to calculate relative cellular accumulation by dividing the slope of the sample molecule by the slope of a reference molecule (Reference mAb#1). All samples were recorded as technical duplicates with three or more experimental replicates (data, statistical analyses and exact number of replicates are provided in Table S1).

### 4. Determination of FcRn-recycling efficiency and relative uptake

To determine relative uptake, the previously described assessment of relative cellular accumulation was used, with the modification of adding a FcRn blocking antibody (re-engineered human IgG1 with the Fv-sequence of SYNT-001 (61)). In short, human primary LSECs were incubated with 200 nM test antibody in the presence of 10 nM FcRn blocking antibody. All samples were recorded as technical duplicates with three or more experimental replicates (data, statistical analyses and exact number of replicates are provided in Table S1). Cells were incubated for 4 h, and further treatment and analysis was carried out as described above. To determine the relative uptake, the normalized MFI_blocked (with FcRn-block) of the antibody formats were divided by the MFI_blocked of the uptake reference antibody (Reference mAb#2). FcRn-recycling efficiencies were calculated from the corrected MFI from the condition with and without FcRn block (4 h timepoint) with the following formula:

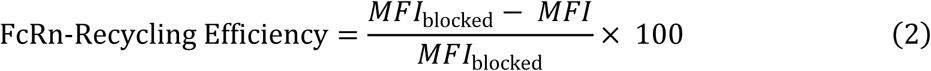

Where MFI_blocked corresponds to the condition with FcRn-block and MFI to the condition without FcRn-block, both for the same antibody format.

### 5. FcRn-affinity chromatography

Analytical FcRn affinity chromatography was performed as previously described (40, 62). Briefly, 30 ug of protein was injected onto a commercial FcRn column (Roche Custom Biotech) at 25 °C with a 0.5 mL/min flow rate. The column was pre-equilibrated with 80 % Buffer A (20 mM MES, 140 mM NaCl, pH 5.5) and 20 % Buffer B (20 mM Tris/HCl, 140 mM NaCl, pH 8.8). Elution was achieved using a 70 minute linear gradient from 20 % to 100 % Buffer B, initiated 10 minutes post-injection. Protein was detected by UV absorbance at 280 nm. To ensure comparability across runs, a standard sample was analyzed at the beginning and end of each sequence and after every 10th run. The relative retention time (RRT) was calculated using the formula:

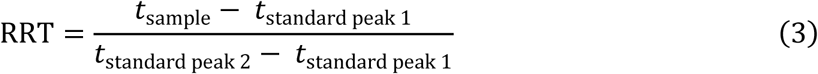

### 6. Pulse-chase FcRn-recycling and transcytosis assay

FcRn-mediated recycling was evaluated using a modified version of the pulse-chase assay (41). Madin-Darby Canine Kidney (MDCK II) cells, engineered to overexpress the alpha-chain of the human neonatal Fc receptor (FcRn) (63), kindly provided by Prof. Walter Hunziker (Singapore University), were cultured as monolayers on Transwell® polycarbonate filters. The assay was performed in a pulse-chase format at 37°C. During the pulse phase, the apical side of the cell monolayer was incubated with the test antibodies at a concentration of 80 nM in a pH 6.0 buffer (HBSS-MES pH 6.0, 0.1 % BSA) for one hour. Following this incubation, the apical and basolateral compartments were washed to remove any unbound antibody. For the chase phase, fresh pH 7.4 buffer (HBSS-HEPES pH7.4, 0.1 % BSA) was added to both compartments, and the cells were incubated for an additional two hours. After the chase period, samples were collected from both the apical (representing recycled antibody) and basolateral (representing transcytosed antibody) compartments. The concentration of the antibody in each sample was quantified using a human IgG ELISA. The total amount of salvaged antibody, referred to as the *total amount of IgG recycled and transcytosed*, was calculated by summing the amounts (ng) of IgG recovered from both the apical and basolateral compartments:

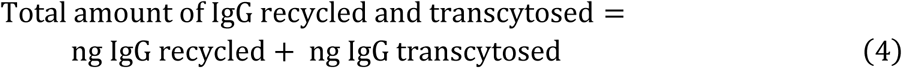

### 7. Heparin chromatography

Heparin chromatography was performed as previously described (21). Briefly, 30 μg of protein in 20 mM histidine buffer (pH 5.5) was injected onto a TSK-Gel Heparin-5PW column (Tosoh Biosciences). Elution was achieved using a 16.5 min linear salt gradient (0-55 % of 1 M NaCl) at 0.8 mL/min, with UV detection at 220 nm. For comparison, relative retention times (RRT) were calculated by normalizing sample retention time to a standard.

### 8. Mouse PK studies and analysis

Mouse single dose PK studies were conducted in hFcRn Tg32 mice (B6.Cg-Fcgrttm1Dcr Tg(FCGRT)32Dcr/DcrJ, Strain #:014565) or in FcRn-KO (specifically, b2m-KO mice, strain B6.129P2-B2mtm1Unc/DcrJ) maintained under ad libitum feeding conditions. All animal related research was conducted in accordance with 2010/63/EU and National legislation regulating the use of laboratory animals in scientific research and for other purposes (Official Gazette 55/13). Single dose pharmacokinetic (SDPK) studies were performed at Selvita d.o.o. (Zagreb, Croatia, on behalf of Roche Pharma Research and Early Development) using microsampling with serial blood collection over 672 hours (Tg32 mice) or 96 (B2m-KO mice). The test antibody was administered as a single intravenous bolus at 3 mg/kg at time 0 h. IVIG (Hizentra) was co-administered intraperitoneally at 1000 mg / kg at two time points: 48 hours prior to mAb administration (t = -48 h) and at 312 h post-mAb administration. Blood samples (0.02 mL) were collected at 0.08, 1.5, 7, 24, 48, 72, 168, 336, 504, and 672 h for Tg32 mice or 0.08, 2, 7, 24, 31, 48, 72, 96 h for b2m-KO mice into serum gel tubes for serum preparation. Each group (antibody study) contained 4 mice.

Collected serum samples were analysed at Microcoat Biotechnologie GmbH (Bernried, Germany, on behalf of Roche Pharma Research and Early Development) using specific ligand binding assays for Binder#1 or golimumab.

For analysis, biotinylated capture reagent specific for the drug antibodies (Roche Diagnostics, Penzberg, Germany) and study samples as well as ruthenylated anti-human-Fc mouse monoclonal antibody (Roche Diagnostics, Penzberg, Germany) and SA-beads, were added stepwise to a detection vessel and incubated for 9 minutes in each step. The SA-beads-bound complex was detected by a measuring cell of a Cobas e411 instrument (Roche Diagnostics, Penzberg, Germany) which counted the SA-beads repeatedly. The quantification of the antibodies was performed by back calculation of the count values using the corresponding calibration curve with a non-linear 4 parameter curve fitting function. All study samples were analyzed in duplicates at one or two dilutions and calibrated against the reference standard. For further data processing and calculation of concentration data the Gen5 software (BioTek Instruments, part of Agilent Technologies, Santa Clara, CA, USA) was used.

PK data were analyzed by noncompartmental analysis using Phoenix WinNonlin (Version 8.5; Certara USA, Inc). In vivo recycling efficiency was calculated by:

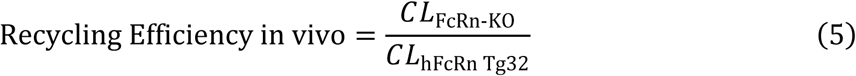

### 9. TNF-α activity assay

The potency of TNF-α and the inhibitory activity of test antibodies were quantified using HEK-Blue-Lucia TNF-α reporter cells (InvivoGen). Cells were maintained in DMEM (4.5 g/l glucose, 2 mM L-glutamine, 10 % (v/v) heat-inactivated FBS (Capricorn), 100 U/ml penicillin-streptomycin (Gibco)) supplemented with 100 ug/ml Normocin and 100 ug/ml Zeocin (InvivoGen). For all assays, cells were harvested and resuspended in identical medium but without Normocin and Zeocin (test medium). Cells were seeded into 96-well plates at 280,000 cells/ml (180 ul/well). To determine the EC80 of TNF-α, a 1:3 serial dilution (starting from 81 ng/ml) was added to the cells. For antibody inhibition assays, 20 ul of test antibody, serially diluted 1:3 in test medium, was added to the cells along with a constant final concentration of 3 ng/ml recombinant human TNF-α (InvivoGen). After overnight incubation (37°C, 5 % CO2), 5 ul of cell-free supernatant was transferred to a 384-well black plate (Corning). Luciferase activity was measured by adding 25 ul of QUANTI-Luc 4 reagent (InvivoGen) and immediately reading the luminescence on a Tecan Infinite M Plex with a 0.1s integration time. All experiments were performed in technical triplicates, and IC50 values were calculated from a 4-parameter logistic curve fit using GraphPad Prism.

### 10. Data analysis, statistical analysis and usage of AI

Data analysis was performed using Microsoft Excel, GraphPad Prism 10, and R4.4.2. Specific software is indicated in the respective methods section. Statistical tests are indicated in the corresponding figure legends and/or the method sections. Details and results of statistical tests are reported along the data in Table S1. No AI tools were used for data analysis and statistical evaluation. Google Gemini was used for reviewing and editing the manuscript.

### 11. Data availability statement

The authors confirm that the data supporting the findings of this study are available within the article and its supplementary materials. Raw datasets are available from the corresponding author upon reasonable request.

## Acknowledgments

The authors would like to thank Alexander Knaupp for his expert advice on the experimental setup of the chromatography methods. We are grateful to Sandra Neuner and Vincent Wieser for their assistance with the data collection of FcRn-and Heparin-affinity chromatography, and to Massimiliano Donzelli for his valuable support during the in vivo studies. We thank Guy Georges for his guidance regarding the design of the FcRn-blocking antibody. Finally, we express our gratitude to Michael Gertz, Jin Zhang, and Martin Lechmann for their critical review and reading of the manuscript. Antibody sketches and schematic method representations were created with BioRender.com

## Author contributions

S.C.W., A.I., H.K., V.H., W.F.R., and T.E.K. designed the research.

S.C.W., J.F., A.-L.B., C.J., S.I.-J., C.S., T.E., T.S. and T.E.K. designed the experiments.

S.C.W., J.F., A.-L.B., A.P., A.C.C. and E.D.L. performed the experiments.

S.C.W., C.J., C.S., A.P., T.E., T.S. and T.E.K. analyzed the data.

S.C.W. and T.E.K. wrote the manuscript.

All authors contributed to and approved the manuscript.

## Supplementary Materials

Supplementary Figures S1–S7

Supplementary Table S1-Data_and_statistical_summary

