## Supplementary Figures for "Spatial architecture of complex antibody formats dictates pharmacokinetics via FcRn-recycling"

Stefan C. Weise *et al.*

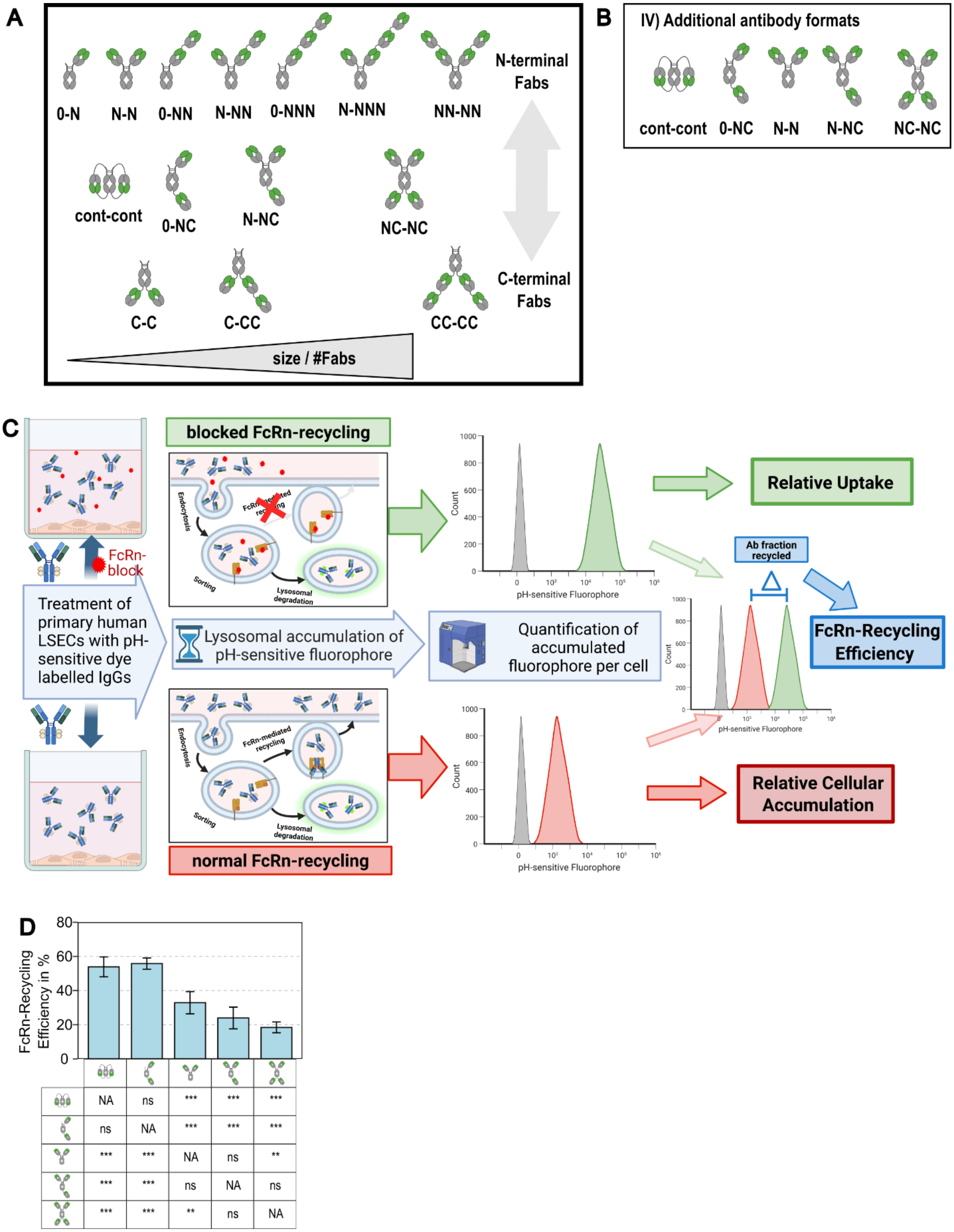


**Figure S1. Binder#1 antibody format design panel, LSEC Assay overview and FcRn-recycling Efficiency of additional antibody formats.**

(**A**) Complete design space of complex antibody formats arranged by molecular weight and N- vs C-terminal Fab arrangement (**B**) Additional antibody formats compared to canonical IgG, not covered in the three design principles (**C**) Schematic overview of the in vitro LSEC assay to determine FcRn-Recycling efficiency, relative uptake and rel. cellular accumulation (rCA). Primary human LSECs were incubated with antibodies labeled with a pH-sensitive fluorophore in the presence or absence of an FcRn-block. During incubation, antibodies in the blocked condition cannot be recycled by FcRn and accumulate in the lysosome, while in the unblocked condition only antibodies which are not rescued by FcRn accumulate in the lysosome. The fluorophore-labeled antibodies accumulated in the lysosome are measured by flow cytometry. The red histogram indicates lysosomal accumulation without the FcRn-block, whereas the green histogram indicates the signal from the FcRn-blocked condition. For calculating the relative uptake, the MFI of the FcRn-blocked condition was normalized to the MFI of a reference antibody. FcRn-Recycling Efficiency was calculated as the percent difference in MFI of the FcRn blocked and FcRn free conditions. rCA was derived from the MFI of the unblocked condition normalized to a reference antibody (details in Method section). (**D**) FcRn-Recycling efficiency of the additional antibody shown in (**B**). Data are represented as mean +/- SD of n=3 - 15 experimental repeats. Statistical analysis was performed using a one-way ANOVA followed by Tukey's multiple comparisons test (ns = not significant, **P < 0.01, ***P < 0.001. Data, exact n and *P* values are provided in Table S1.

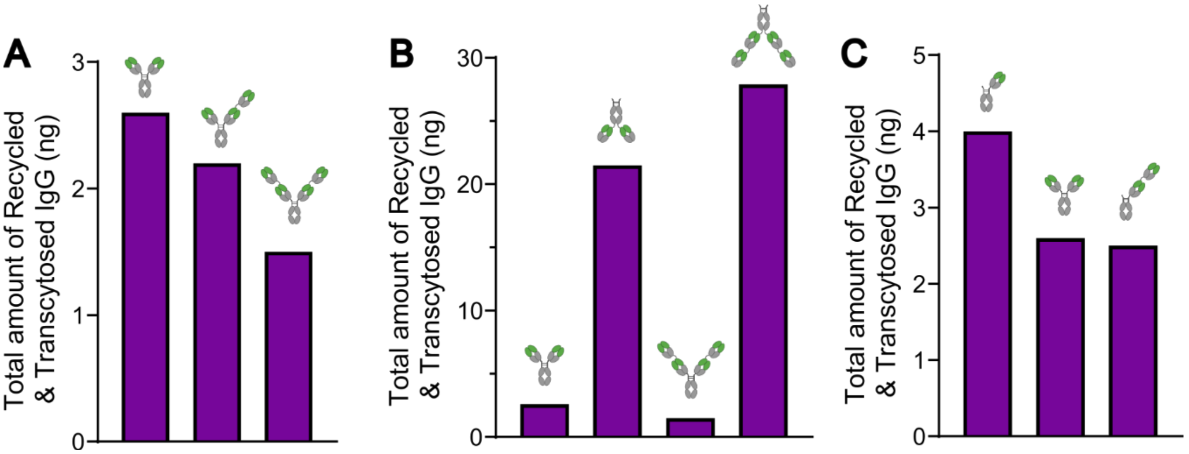


**Figure S2. Results of the Pulse-Chase FcRn-recycling assay confirm the impact of antibody format.** (**A**–**C**) FcRn-mediated recycling and transcytosis measured in vitro by a pulse-chase experiment. Representative antibody formats with Binder#1 were chosen for (**A**) increasing number of Fabs, (**B**) C- compared to N-terminal Fab location and (**C**) one- compared to dual armed antibodies.

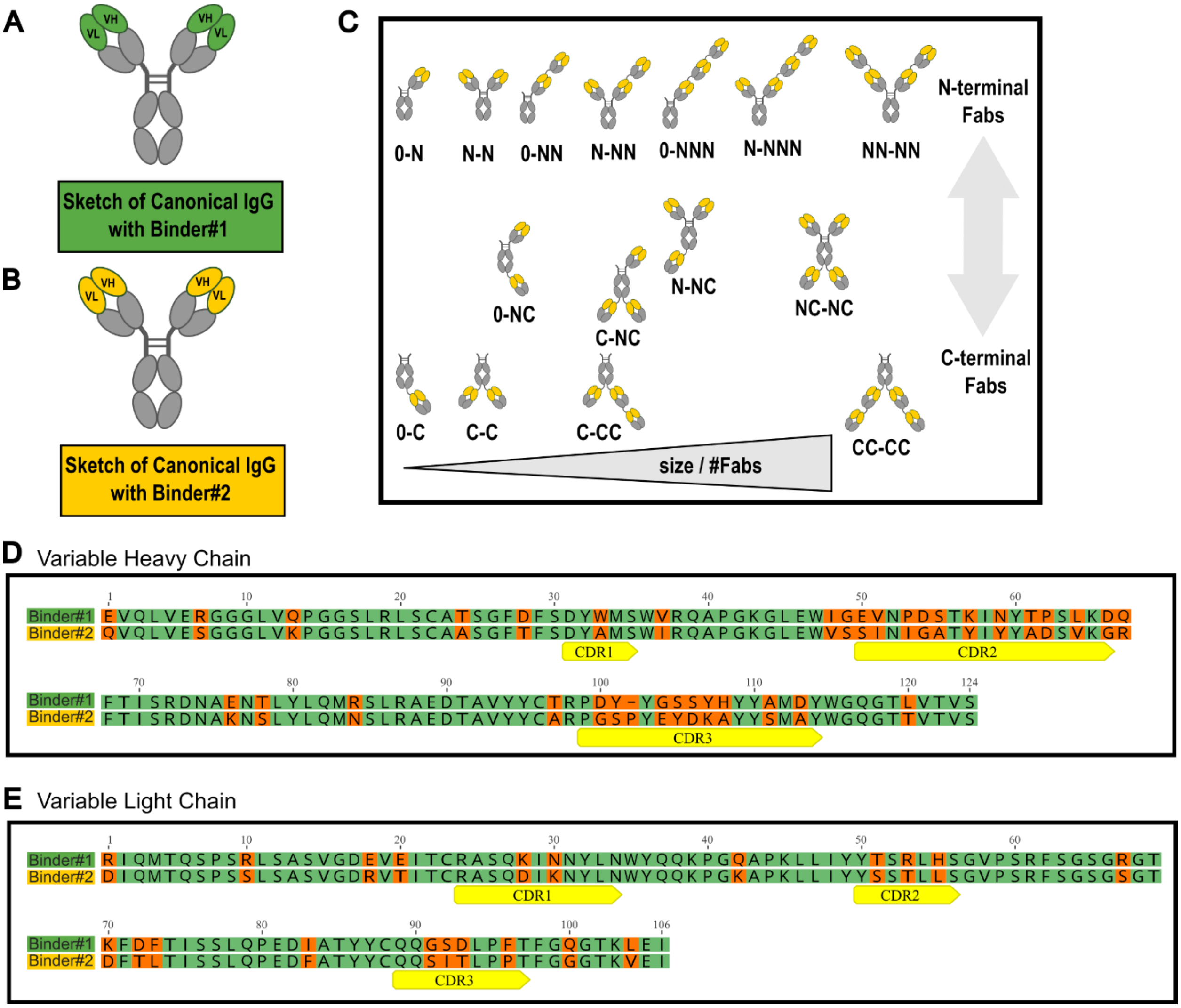


**Figure S3. Comparison of the Fv-sequences between Binder#1 and #2.** Sketch of a canonical IgG with Binder#1 (**A**) and Binder#2 (**B**). (**C**) Schematic representation of the complete set of complex antibody formats produced with Binder#2. Fabs containing Binder#2 are indicated in yellow. (**D** and **E**) Amino acid sequence alignment of the (**D**) variable heavy (VH) and (**E**) variable light (VL) chains indicating differences between Binder#1 and Binder#2 sequences. CDRs are highlighted in yellow.

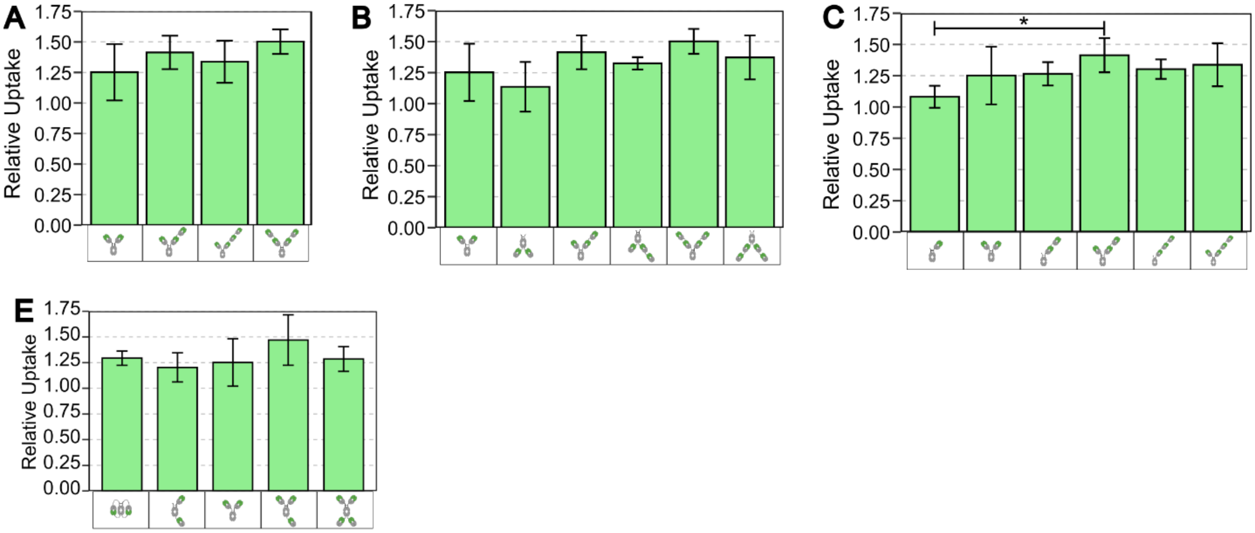


**Figure S4. Binder#1 antibody formats show similar relative uptake** (**A**–**D**) Bar graphs representing relative uptake of Binder#1-formats grouped by: increasing N-terminal number of Fabs (**A**), comparison of C-terminal Fab location to N-terminal (**B**), comparing one- against dual-armed antibodies (**C**) and additional formats (**D**). Represented are mean +/- SD of n=3-15 independent experiments. Statistical analysis was performed using a one-way ANOVA [$F(13, 66) = 2.67$, $P = 0.0047$] followed by Tukey's multiple comparisons test. *: $P < 0.05$. Data, exact n and $P$ values are provided in Table S1.

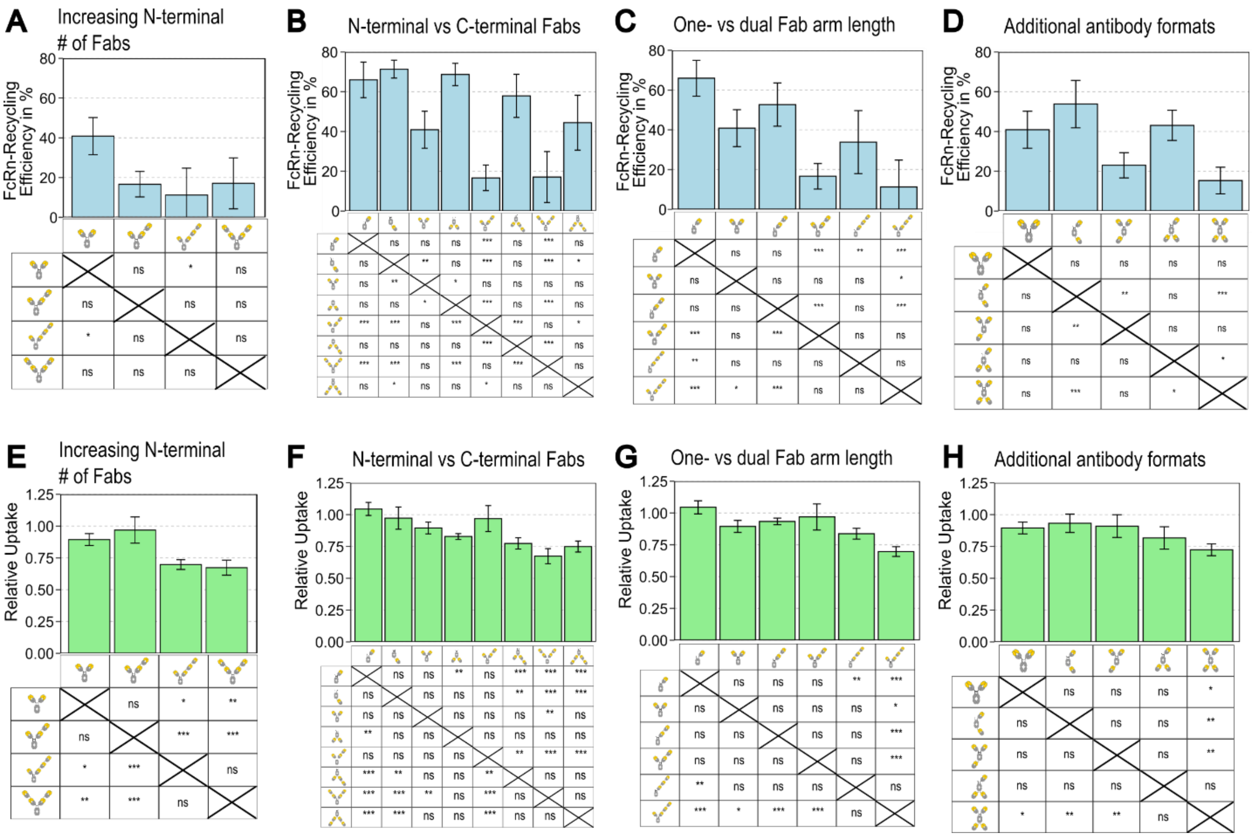


**Figure S5. Binder#2 antibody formats exhibit format-dependent trends in FcRn-recycling efficiency and relative uptake** Bar graphs representing FcRn-recycling efficiencies (**A–D**) and relative uptake (**E–H**) of Binder#2-formats combinations grouped by: increasing N-terminal number of Fabs (**A** and **E**), comparison of C-terminal Fab location to N-terminal (**B** and **F**), comparing one- against dual-armed antibodies (**C** and **G**) and additional formats (**D** and **H**). Represented are mean +/- SD of n = 3-4 experimental repeats. Statistical analysis was performed using a one-way ANOVA [FcRn-recycling efficiency: $F(14, 43) = 15.82$, $P < 0.0001$; Relative Uptake: $F(14, 43) = 11.75$, $P < 0.0001$] followed by Tukey's multiple comparisons test. ns = not significant, *: $P < 0.05$, **: $P < 0.01$, ***: $P < 0.001$. Data, exact n and $P$ values are provided in Table S1.

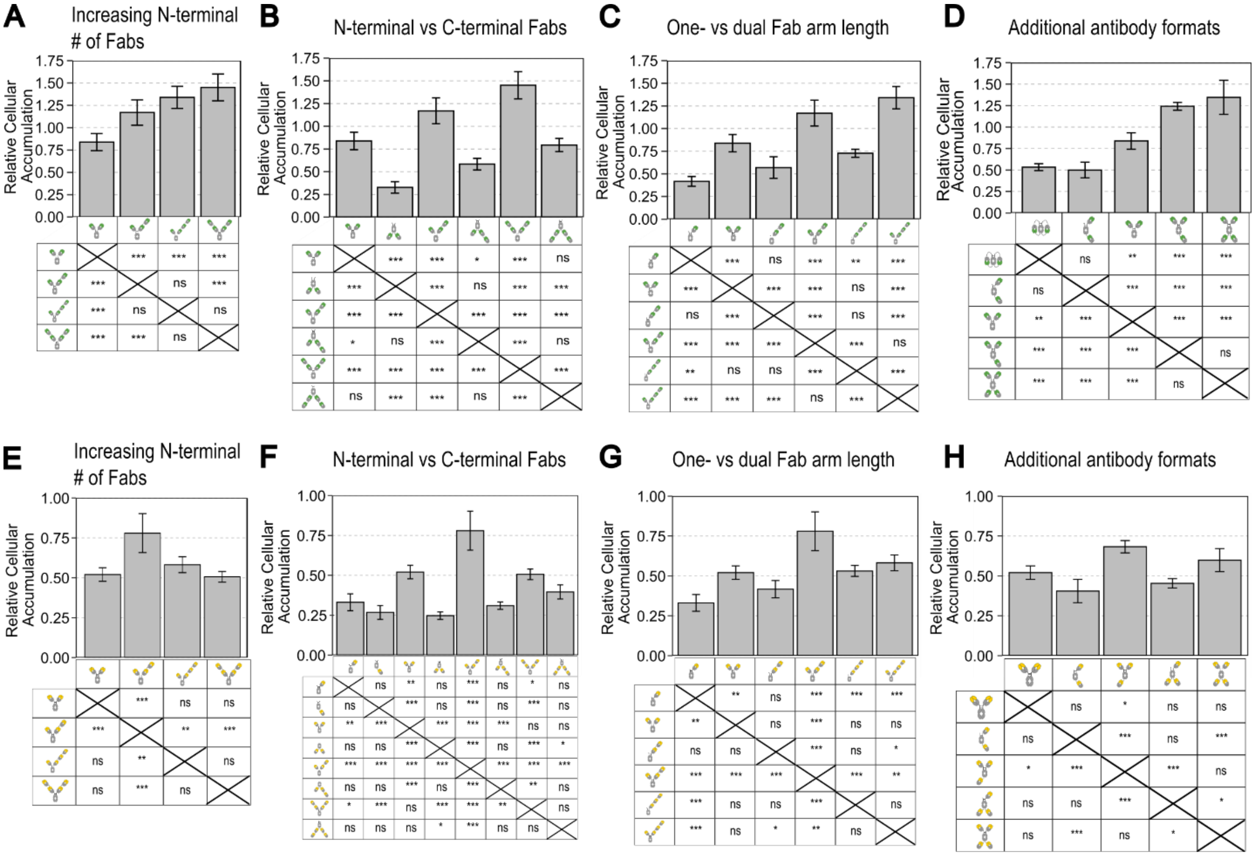


**Figure S6. Binder#1 and #2 antibody formats show significant differences in relative cellular accumulation** Bar graphs showing the relative cellular accumulation (rCA) of Binder#1-formats combinations (**A–D**) and Binder#2-formats combinations (**E–H**) grouped by increasing N-terminal number of Fabs (**A** and **E**), comparison of C-terminal Fab location to N-terminal (**B** and **F**), comparing one- against dual-armed antibodies (**C** and **G**) and additional formats (**D** and **H**). Represented are mean +/- SD of n = 3-19 experimental repeats. Statistical analysis was performed using a one-way ANOVA. (**E–H**)**:** [$F(14, 43) = 30.18$, $P < 0.0001$], (**A–D**)**:** [$F(13, 72) = 64.07$, $P < 0.0001$] followed by Tukey's multiple comparisons test. ns = not significant, *$P < 0.05$, **$P < 0.01$, ***$P < 0.001$. Data, exact n and *P* values are provided in Table S1.

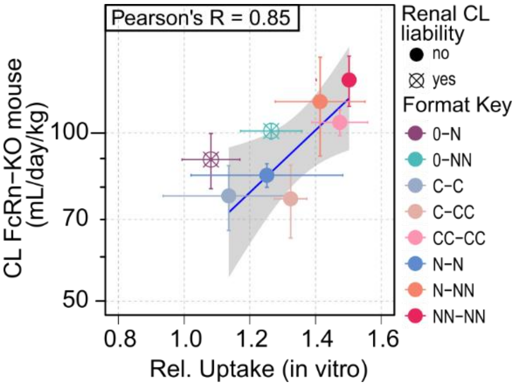


**Figure S7. CL of the Binder#1 antibody formats in FcRn-KO mice correlates to the relative uptake measured in the LSEC assay** Plot depicting the correlation between relative uptake (in vitro) and clearance in the FcRn-KO model. Single-armed antibody formats indicated with renal CL liability were excluded from the regression and correlation analysis. Blue line and grey area show the linear regression with the 95% confidence interval. Pearson correlation between relative uptake (in vitro) and clearance in the FcRn-KO model (R = 0.85, P = 0.031, n = 6).
